# PAK6 rescues pathogenic LRRK2-mediated ciliogenesis and centrosomal cohesion defects in a mutation-specific manner

**DOI:** 10.1101/2024.04.11.589075

**Authors:** Lucia Iannotta, Rachel Fasiczka, Giulia Favetta, Yibo Zhao, Elena Giusto, Elena Dall’Ara, Jianning Wei, Franz Y. Ho, Claudia Ciriani, Susanna Cogo, Isabella Tessari, Ciro Iaccarino, Maxime Liberelle, Luigi Bubacco, Jean-Marc Taymans, Claudia Manzoni, Arjan Kortholt, Laura Civiero, Sabine Hilfiker, Michael Lu, Elisa Greggio

## Abstract

P21 activated kinase 6 (PAK6) is a serine-threonine kinase with physiological expression enriched in the brain and overexpressed in a number of human tumors. While the role of PAK6 in cancer cells has been extensively investigated, the physiological function of the kinase in the context of brain cells is poorly understood. Our previous work uncovered a link between PAK6 and the Parkinson’s disease (PD)-associated kinase LRRK2, with PAK6 controlling LRRK2 activity and subcellular localization *via* phosphorylation of 14-3-3 proteins.

Here, to gain more insights into PAK6 physiological function, we performed protein-protein interaction arrays and identified a subgroup of PAK6 binders related to ciliogenesis. We confirmed that endogenous PAK6 localizes at both the centrosome and the cilium, and positively regulates ciliogenesis not only in tumor cells but also in neurons and astrocytes. Strikingly, PAK6 rescues ciliogenesis and centrosomal cohesion defects associated with the G2019S but not the R1441C LRRK2 PD mutation. Since PAK6 binds LRRK2 via its GTPase/Roc-COR domain and the R1441C mutation is located in the Roc domain, we used microscale thermophoresis and AlphaFold2-based computational analysis to demonstrate that PD mutations in LRRK2 affecting the Roc-COR structure substantially decrease PAK6 affinity, providing a rationale for the differential protective effect of PAK6 toward the distinct forms of mutant LRRK2.

Altogether, our study discloses a novel role of PAK6 in ciliogenesis and points to PAK6 as the first LRRK2 modifier with PD mutation-specificity.

## Introduction

P21-activated kinases (PAKs) comprise a group of serine-threonine kinases that cover crucial roles in signal transduction events. They exert their activity downstream of Rho GTPases by modulating the cytoskeleton, facilitating gene transcription, and promoting cell survival (1). In mammals, the family is divided in two groups, namely group I PAKs (PAK1, 2 and 3) and group II PAKs (PAK4, 5 and 6). All PAKs are characterized by a highly conserved kinase domain at the C-terminus, whereas the N-terminal regulatory regions are more divergent but all contain a Cdc42 and Rac-interactive binding (CRIB) domain required for activation (group I) or relocalization (group II) of the kinases (2,3). Among group II PAKs, PAK6, along with its homologue PAK5, is highly expressed in neurons (4) and, accordingly, Pak5/Pak6 double knockout mice exhibit locomotor impairment and learning and memory defects (5,6). Of note, PAK6 expression is upregulated in a number of cancers (7–10) and PAK6 interaction with androgen receptor (AR) promotes its phosphorylation and prostate cancer cell motility and invasion (11). A genome-wide coexpression analysis of steroid receptors in the mouse brain identified PAK6 as a mediator for the effects of AR on dopaminergic transmission, suggesting the relevance of PAK6 activity in dopaminergic neuron physiology (12).

Parkinson’s disease (PD) is a neurodegenerative movement disorder mainly affecting the dopaminergic nigrostriatal pathway (13). Despite being predominantly sporadic, PD can manifest with a hereditary pattern due to mutations in risk factor or causal genes (14). Mutations in *Leucine-rich repeat kinase 2* (*LRRK2*) represent a common cause of familial PD (15). *LRRK2* encodes a large multidomain protein equipped with a GTPase Roc-COR domain, a serine-threonine kinase domain and a number of protein-protein interaction domains (16). Mechanistically, LRRK2 mutations increase kinase activity by enhancing LRRK2 substrate phosphorylation through different pathways: pathological mutations localized in the Roc-COR domain (e.g. R1441C/G/H and Y1699C) favor the GTP-bound state and/or decreased 14-3-3 protein binding, thus promoting LRRK2 localization at membranes where its Rab GTPase substrates are localized (17,18). Conversely, the G2019S mutation in the kinase domain shifts the equilibrium from the inactive to the active state, thus accelerating substrate phosphorylation (18). LRRK2 phosphorylates a subset of Rab GTPases, including Rab3, Rab8a, Rab10, Rab12 and Rab35 (19), to control lysosomal stress response (20,21), phagocytosis (22,23) and primary cilia formation (24). In the presence of pathogenic G2019S and R1441C/G LRRK2 mutations, Rab8a/Rab10 hyperphosphorylation results in impairment of primary cilia formation in neurons and astrocytes (25–27) as well as centrosome cohesion defects (28–31). The primary cilium serves as an antenna-like organelle, facilitating the transmission of external signals to the nucleus to uphold cellular homeostasis. Anchored at the centrosome, it extends from the cell surface in close proximity to the nucleus (32).

Our previous work has provided multiple lines of evidence for a bidirectional connection between LRRK2 and PAK6. For example, PAK6 and LRRK2 interact through their CRIB and Roc-COR domains, respectively (33), and this interaction is required for PAK6 to control neurite outgrowth in the mouse striatum (33). In addition, PAK6 kinase activity can negatively regulate LRRK2 phosphorylation at Ser910 and Ser935 through phosphorylation of 14-3-3 proteins (34), with consequent reduction of LRRK2 kinase activity (35,36). Moreover, PAK6 effectively reduces Rab10 phosphorylation upon expression of mutant LRRK2 G2019S (35) and rescues the G2019S LRRK2-associated neurite shortening phenotype (34), but it fails to do so in the presence of the Roc-COR R1441G LRRK2 mutation (35). Of note, patients affected by sporadic and G2019S LRRK2-linked PD exhibit changes in the amount of circulating PAK6 and of its substrate and LRRK2 interactor 14-3-3γ (37), highlighting the potential clinical relevance of this pathway.

Here, starting from unbiased analyses of PAK6 interactomes, we observed that PAK6 binds proteins associated with the primary cilium and confirmed its localization at this organelle. We further established PAK6 as a positive regulator of ciliogenesis in different cell types, including primary neurons and astrocytes. Importantly, we found that PAK6 rescues G2019S LRRK2-mediated ciliogenesis and centrosomal cohesion defects but it is unable to provide protection under R1441C LRRK2 expression. By combining microscale electrophoresis (MST) analysis and alpha-fold modelling, we demonstrated that the binding of the CRIB domain to Roc-COR is weakened by pathological substitutions at R1441 and Y1699 LRRK2 residues, which are both sitting at the binding interface, providing a rationale for the observed PD mutation-specific effect exerted by PAK6.

## Results

### PAK6 interacts with primary cilium proteins

To gain insights into the physiological function of PAK6, we undertook an unbiased approach to identify novel protein-protein interaction (PPI) partners. A Human Proteome Microarray, containing more than 20,000 recombinant proteins, was incubated with recombinant full-length human PAK6, leading to the identification of several candidate interactors (**Figure 1a**). To search for common biological pathways, we performed a gene ontology (GO) analysis of hits with Z score > 2.5 (147 proteins) (**Supplemental table 1**). A term size cutoff of 2000 was applied in order to increase specificity. Significant gene ontology biological process (GO:BP) terms were manually grouped into semantic categories to reduce redundancy, leading to processes related to catabolism, cytoskeletal dynamics, protein modification, response to hormones, muscle contraction, and cell projection organization (**Figure 1b**). In parallel, we retrieved PAK6 PPI using the online PPI query tools PINOT, HIPPIE and MIST (hereafter PHM) (38) to compare our protein array findings with already available PPI datasets from the literature (**Supplemental table 1**). Similarly, PPI were subjected to GO:BP analysis with 1000 term size cutoff and grouping into semantic categories. Across the 2 lists, we found both different and overlapping biological processes, which is expected given that PHM PPI validated interactors have been identified with complementary experimental approaches (AP/MS; co-IP; yeast-2-hydrid). Overlapping categories included response to stimuli/hormones and cell projection organization (**Figure 1b)**. Consistent with the involvement of PAK6 in cell projection processes, we noted that a top candidate in the PAK6 array was CLUAP1 (**Figure 1a**), an evolutionary conserved protein promoting ciliogenesis (39,40). Thus, we subsequently crossed the primary cilium proteome (GO:0005929, 640 genes) with either the PAK6 array PPI dataset or the PAK6 PHM PPI dataset, and found 6 (CLUAP1, ENTR1, AK7, CCDC40, DRC3 and SUFU) and 3 (AKT1, APP, PKM) overlapping proteins, respectively (**Figure 1c**). In accordance, Gene Set Enrichment Analysis (GSEA) results suggested that both of the lists were significantly enriched for ciliary proteins (*P* values = 0.036 for the PAK6 PHM PPI dataset and 0.041 for the PAK6 array PPI dataset), identifying 6 (CLUAP1, ENTR1, AK7, CCDC40, DRC3 and SUFU) and 3 (AKT1, APP, PKM) cilium-related proteins, respectively (**Figure 1c**, **Supplemental table 2)**. Together with PAK6, these 10 proteins formed a physical and functional network (**Figure 1d**), supporting the notion that they are at least partially biologically connected as a group (number of nodes: 10; number of edges: 8; expected number of edges: 3; PPI enrichment *P* value = 0.014). Altogether, these results point to PAK6 as a novel player in ciliogenesis-related pathways.

**Figure 1.**
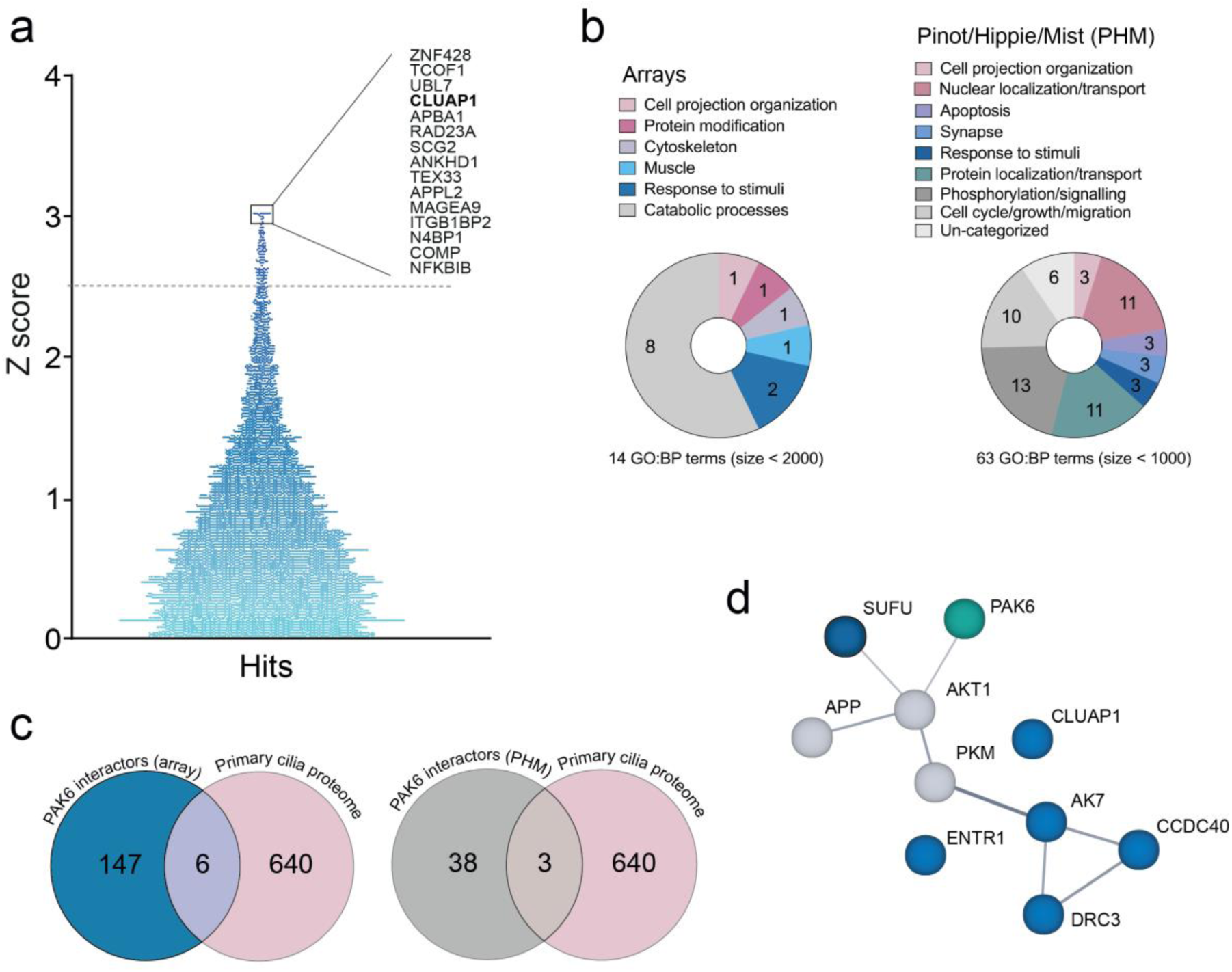
PAK6 interacts with ciliary proteins. **(a)** Distribution of PAK6 candidate interactors according to their Z-score retrieved from a Human Proteome Microarray probed with recombinant full-length human PAK6. **(b)** A GO:BP analysis using gProfiler g:GOSt (https://biit.cs.ut.ee/gprofiler/gost) was performed for PAK6 candidate interactors with Z score >2.5 (left) and for PAK6 interactors annotated in PPI web-based tools PINOT, HIPPIE and MIST (PHM) (right). GO:BP terms with 2000 (array) and 1000 (PHM) term size were grouped into semantic categories. **(c)** Venn diagrams showing overlaps between the primary cilium proteome (GO:0005929, 640 genes) and the experimental (array) PAK6 interactome (left) or the literature-based (PHM) PAK6 interactome (right). **(d)** Protein network of overlapping PAK6 interactors with the primary cilium proteome **(c)** (including PAK6) obtained with STRING (https://string-db.org/cgi/input?sessionId=b1S4T5BW27rz&input_page_show_search=on); number of nodes: 11, number of edges: 11, average node degree: 2, average local clustering coefficient: 0.591, expected number of edges: 3, PPI enrichment *P*-value: 0.000502. Blue nodes are ciliary proteins present in the experimental PAK6 interactome (array) and grey nodes are those found in the literature-based PAK6 interactome.

### PAK6 is localized at centrosomes and primary cilium and positively regulates ciliogenesis

Using an anti-PAK6 antibody that recognizes the native PAK6 protein, we investigated the subcellular localization of PAK6 in different cell types, including mouse embryonic fibroblasts (MEFs), breast cancer MCF7 cells and HEK293T cells. In wild-type MEF cells, a portion of Pak6 co-localizes with the centrosome marker γ-tubulin, with signal specificity confirmed in MEFs derived from Pak6 null mice (**Figure 2a**). PAK6-centrosome association was also confirmed in MCF7 cells by co-localization with γ-tubulin (**Figure 2b**) and biochemically by co-fractionation of PAK6 with γ-tubulin in centrosome-enriched subcellular fractions using a sucrose gradient (**Figure 2c**). The localization of PAK6 at the centrosome prompted us to further determine its relationship with primary cilia. Co-staining of HEK293T cells against the centrosomal marker pericentrin and the primary cilium marker Arl13b confirmed PAK6 localization at the centrosome/basal body and highlights that the kinase is also localized within the primary cilium axoneme (**Figure 2d**). To investigate whether PAK6 localization at the centrosome/cilium impacts cilia biology, we downregulated PAK6 by lentivirus-mediated shRNA expression in HEK293T cells. Seventy-two hours post-transduction, primary cilia were induced by serum free media overnight. The control scrambled shRNA infected HEK293T cells responded to serum starvation with robust ciliogenesis. However, shRNA-mediated PAK6 knock-down significantly downregulated both cilia number and length (**Figure 2e**). Similarly, MEFs isolated from Pak6 KO mice display reduced cilia length and number (**Figure 2f**). Taken together, these data support our PPI screens of PAK6 interacting with ciliary proteins.

**Figure 2.**
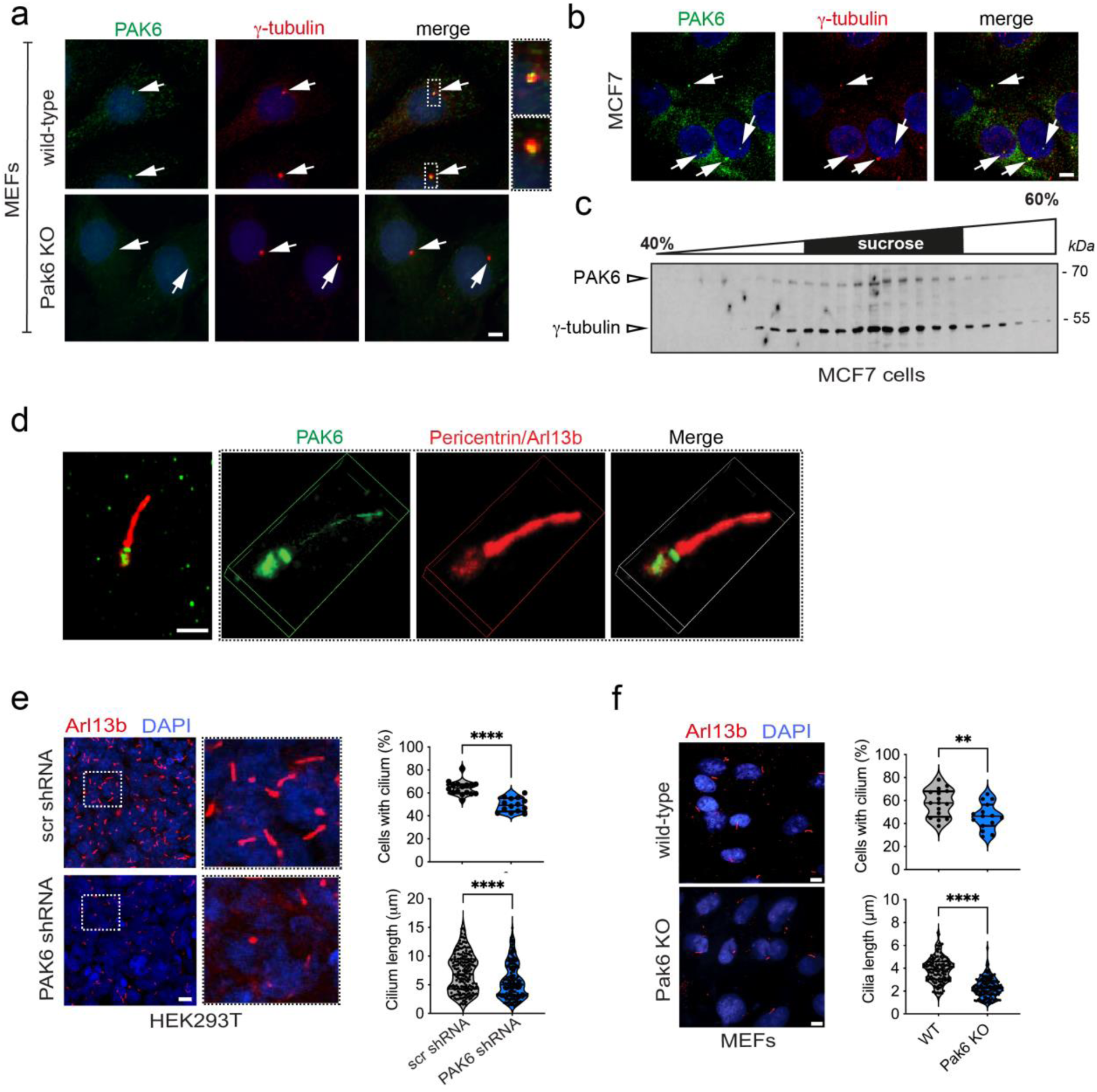
PAK6 is localized at centrosomes and primary cilium and regulates ciliogenesis. **(a)** Localization of Pak6 at centrosome. Staining of mouse embryonic fibroblasts (MEFs) derived from either wild type (MEFs WT) mice or Pak6 null (MEFs KO) mice with anti-γ-tubulin (mouse) and anti-PAK6 (rabbit) antibodies. Insets show localization of Pak6 at centrosome. Scale bar = 10 μm. **(b)** Localization of PAK6 at centrosome in MCF7 cells as evidenced by immunostaining with anti-γ-tubulin (mouse) and anti-PAK6 (rabbit) antibodies. **(c)** Co-sedimentation of PAK6 with γ-tubulin in sucrose gradient fractions. Cell lysates from MCF7 cells were subjected to 40-60% sucrose density gradient ultra-centrifugation and fractionated. The resulting fractions were resolved by SDS-PAGE and analyzed by immunoblot with anti-PAK6 and anti-γ-tubulin antibodies. **(d)** PAK6 localizes to the basal body of primary cilium. Immuno-staining of PAK6 (green) localizes it to the basal body of the primary cilium in HEK293T cells. The primary cilium axoneme and basal body were respectively identified by staining with Arl13b (red) and anti-pericentrin (red). Scale bar = 2 μm. **(e)** PAK6 regulates ciliogenesis in HEK293T cells. Knock-down of PAK6 expression by PAK6 specific lentiviral-mediated shRNA downregulates ciliogenesis as shown by a decrease in the percentage of ciliated cells and a decrease in ciliary length as compared to scrambled shRNA. Cilia were stained with antibodies against Arl13b. Violin plots represent the percentage of cells per field with a primary cilium (top; N=3 experiments: n(scramble)=16, n(Pak6 shRNA)=15 fields) and the length of cilia (bottom; n(scramble)=191, n(Pak6 shRNA)=200 cilia). Unpaired t-test, \*\*\*\**P* < 0.0001. Scale bar = 10 μm. **(f)** Pak6 regulates ciliogenesis in MEF cells. Pak6 KO MEF cells also exhibit a deficit in the percentage of ciliated cells and ciliary length. Cilia were stained with antibodies against Arl13b. Violin plots represent the percentage of cells per field with a primary cilium (top; N=3 experiments: n(WT)=16, n(Pak6 15)= fields) and the length of cilia (bottom; n(WT)=142, n(Pak6 KO)=150 cilia). Unpaired t-test, \*\*\*\**P* < 0.0001. Scale bar = 10 μm.

### PAK6 promotes ciliogenesis in brain cells

While being overexpressed in a number of cancers, under physiological conditions PAK6 shows a restricted tissue expression with enrichment in the brain. This can be inferred from RNA expression datasets (https://www.proteinatlas.org/) and co-expression with genes involved in neuronal/synaptic functions (https://seek.princeton.edu/seek/; (41). In particular, significant GO:BP terms (with electronic annotations) enriched from SEEK co-expression analysis belong to semantic categories related to neuron development, synaptic transmission and plasticity, response to signals, astrocyte differentiation and, consistently, cell projection organization (**Supplemental Table 3** and **Figure 3a**). To investigate the involvement of PAK6 in ciliogenesis in brain cells, we generated polyclonal stable neuroblastoma SH-SY5Y cells overexpressing PAK6 via lentiviral vector (PAK6 OE) and, as control, downregulated PAK6-OE cells with LV-shRNA against PAK6 (PAK6 OE + PAK6 shRNA) (**Figure 3b**). Western blot analysis confirmed efficient overexpression of PAK6 and almost complete downregulation by shRNA. Moreover, PAK6 is active as evidenced by the presence of phospho-S560, a PAK6 autophosphorylation site sitting in a conserved motif shared with PAK4 and PAK5 (**Figure 3b**).

**Figure 3.**
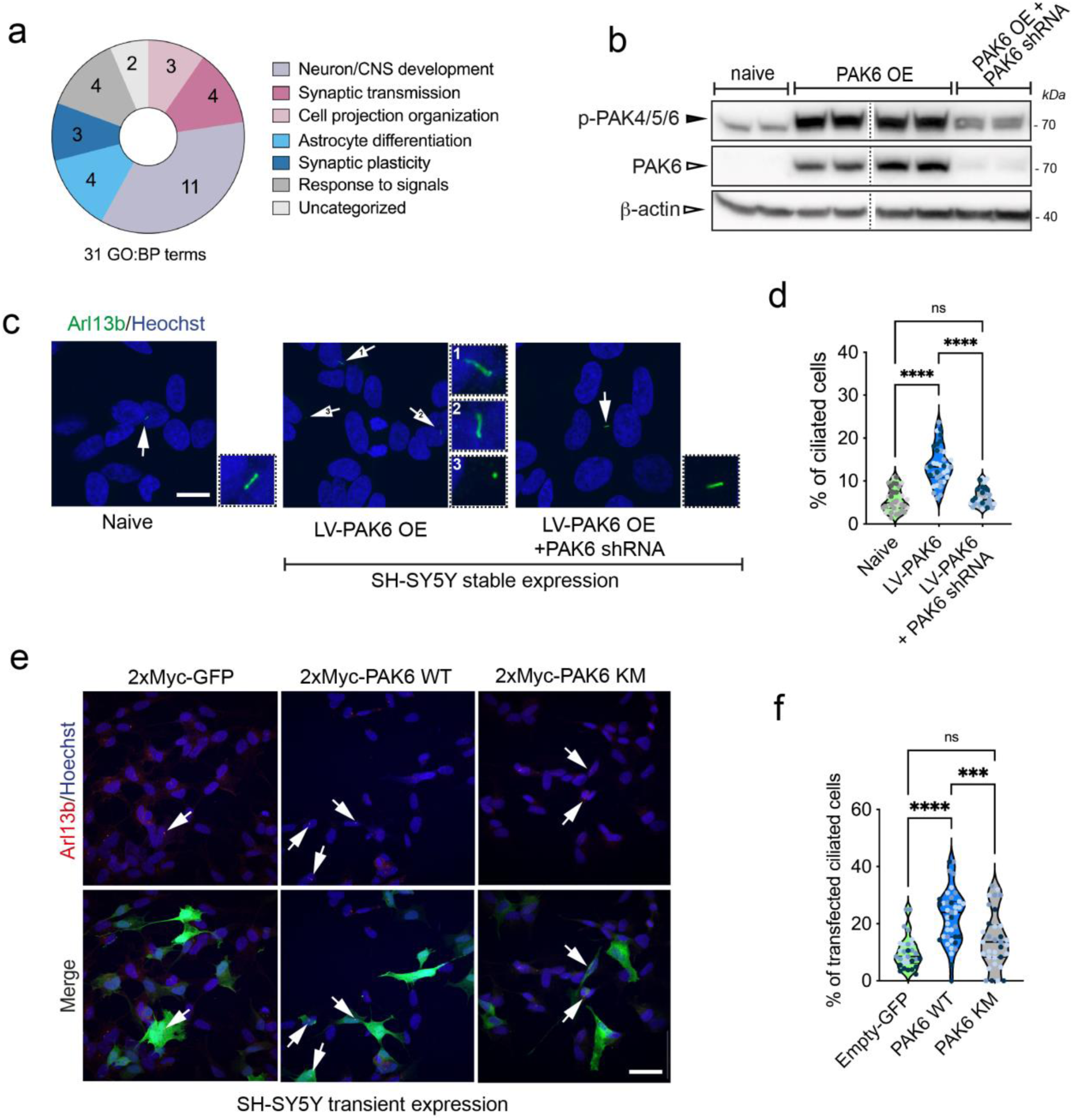
PAK6 promotes ciliogenesis in SH-SY5Y cells. **(a)** Semantic categories from GO:BP (electronic annotation) PAK6 co-expression analysis using Search-Based Exploration of Expression Compendium (SEEK) (41). **(b)** Western blot analysis of PAK6 and phospho-PAK6 in naïve, stable LV-PAK6 and stable LV-PAK6 shRNA SH-SY5Y cells. **(c)** Example of SH-SY5Y naïve, stable LV-PAK6 and stable LV-PAK6 shRNA SH-SY5Y cells. Cilia were stained anti-Arl13b (green) and nuclei with DAPI. Arrows point to primary cilia. Scale bar 10 μm. **(d)** Quantification of (c). Violin plots represent the percentage of cells per field with a primary cilium. N=3 experiments; n(naïve)=37, n(LV-PAK6 OE)=36, n(shRNA PAK6)= 31 fields analyzed; one-way ANOVA with Tukey’s post-test (\*\*\*\**P* < 0.0001). **(e)** Example of SH-SY5Y transfected with 2xMyc-GFP, 2xMyc-PAK6 WT and 2xMyc-PAK6 K436M (KM). Cilia were stained anti-Arl13b (red), and GFP or PAK6 with anti-Myc (green) antibodies, and nuclei with DAPI. Arrows point to primary cilia in transfected cells. Scale bar 20 μm. **(f)** Quantification of (e). Violin plots represent the percentage of transfected cells per field with a primary cilium. N=3 experiments; n(GFP)=30, n(PAK6 WT)=39, n(PAK6 KM)= 33 fields analyzed; one-way ANOVA with Tukey’s post-test (\*\*\*\**P* < 0.0001; \*\*\**P* < 0.001).

Next, we compared the number of ciliated cells across naïve, PAK6 OE and PAK6 OE+PAK6 shRNA using Arl13b staining. The percentage of ciliated cells (low in the absence of serum starvation in non-differentiated SH-SY5Y cells) was 5.6 ± 0.5 (mean ± SEM, n=35 cells, N=3) in naïve SH-SY5Y, whereas in PAK6 OE cells the percentage was increased to 13.7 ± 0.7 (n=36 cells, N=3). Importantly, downregulation of PAK6 in PAK6 OE returned the number of ciliated cells to the control level (6.3% ± 0.4, n=31 cells, N=3). Similarly, the length of the cilium was increased in PAK6 OE cells and returned to the control (naïve) level in PAK6 OE+PAK6 shRNA cells (naïve: 1.48 ± 0.07 µm, PAK6 OE: 1.90 ± 0.04 µm, PAK6 OE+PAK6 shRNA: 1.6 ± 0.06 µm), overall supporting the notion that PAK6 acts as a positive regulator of ciliogenesis in SH-SY5Y cells.

To complement these data and explore the effect of PAK6 kinase activity, we transiently overexpressed 2xMyc-PAK6 wild-type (WT) and 2xMyc-PAK6 K436M (KM), kinase dead (34) along with 2xMyc-GFP control in SH-SY5Y naïve cells, stained for the cilia marker Arl13b and counted the number of transfected SH-SY5Y cells that were ciliated. As shown in **figure 3e-f**, PAK6 WT increased the number of ciliated cells compared to control, while the effect of kinase dead PAK6 KM was not statistically significant. Overall, these experiments indicate that 1) PAK6 is a positive regulator of ciliogenesis and 2) PAK6 kinase activity is required to promote this process.

Next, to translate these findings to more physiologically relevant cellular models, we isolated primary cortical neurons from Pak5/Pak6 double knock-out (dKO) mice (5) and stained for the neuronal ciliary marker AC3 after 14 days *in vitro*. We determined both ciliary length and frequency (ratio of ciliated cells) since a change in either one of those parameters will affect appropriate ciliary signaling. Both WT and Pak5/Pak6 dKO cultures displayed a similar number of ciliated cells (∼60%), however the morphology was different between genotypes, with Pak5/Pak6 dKO cilia being shorter than WT cilia (∼3 µm vs ∼3.5 µm) (**Figure 4a-b**). A shorter primary cilium in Pak5/Pak6 null neurons suggests that the stability of this organelle is affected in the absence of these kinases. Since PAK6 is co-expressed with genes involved in astrocyte differentiation (**Figure 3a**), we next isolated primary striatal astrocytes from WT and Pak5/Pak6 dKO mice and stained with Arl13b antibodies. The number of ciliated astrocytes was lower in Pak5/Pak6 dKO astrocytes (∼40% vs ∼25%) while the length remained unaltered (**Figure 4c-d**). To rule out that the effect observed was mediated by Pak5 or by a combination of Pak5 and Pak6 activities, 3xFlag-PAK6 WT or 3xFlag-GUS control were ectopically re-expressed in the Pak5/Pak6 dKO null background and the proportion of transfected ciliated cells counted. As illustrated in **figure 4e-f**, PAK6 expression was sufficient to fully rescue the reduced cilia number of Pak5/Pak6 dKO astrocytes (3xFlag-GUS ∼ 26%; 3xFlag-PAK6 ∼40%). Altogether, these data support PAK6 as a positive regulator of ciliogenesis in both mouse primary neurons and astrocytes.

**Figure 4.**
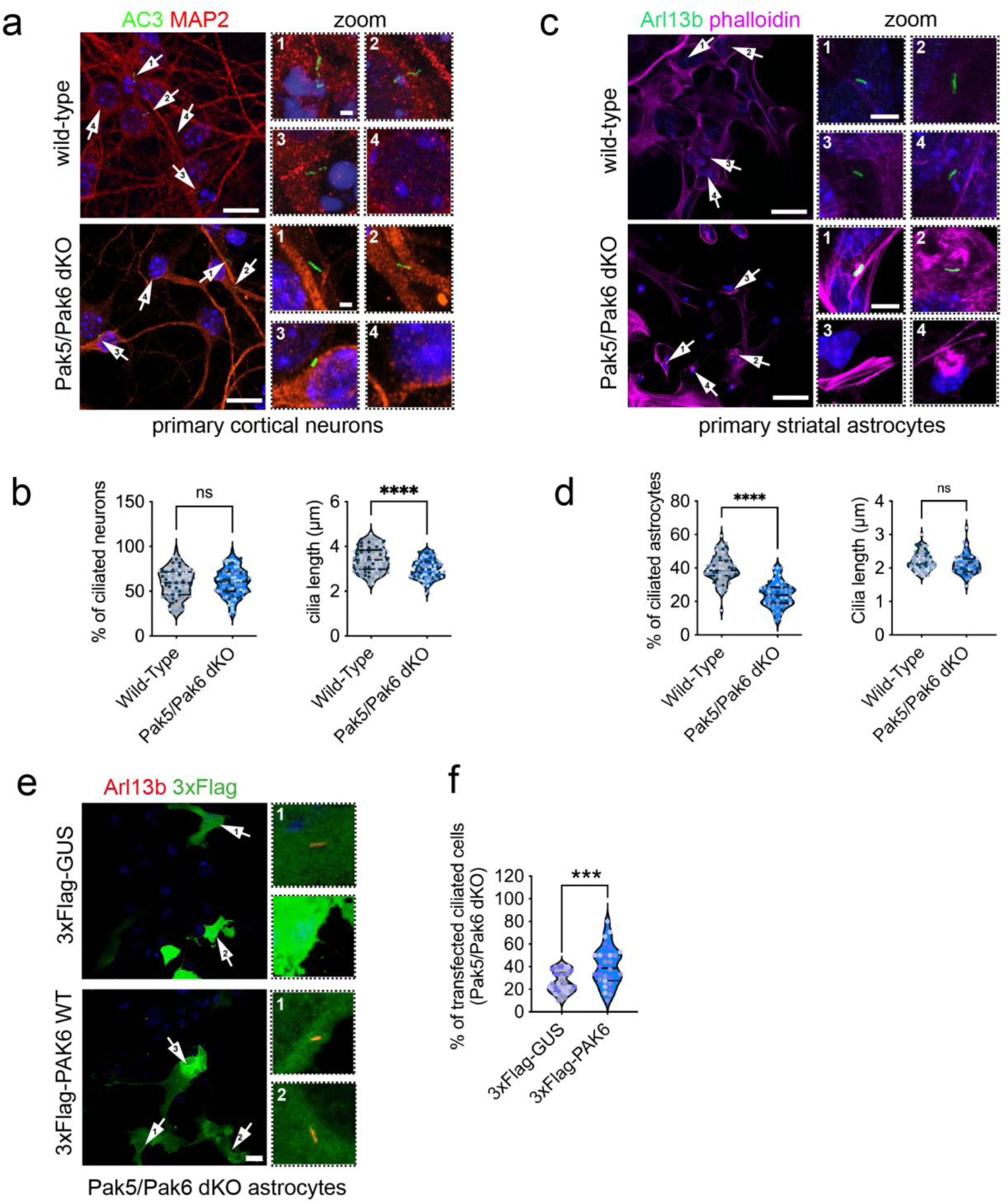
PAK6 promotes ciliogenesis in primary neurons and astrocytes. **(a)** Example of primary cortical neurons isolated form Pak5/Pak6 dKO mice. Immunocytochemistry was performed with antibodies anti-AC3 (neuronal cilia marker, green), anti-MAP2 (neuronal marker, red) and with DAPI (blue). Arrows point to primary cilia (scale bar 10 µm). Scale bar of zoomed images is 1 µm. **(b)** Quantification of (a). Violin plots represent the percentage of cells per field with a primary cilium (left) and the cilia length (right). N=3 experiments; n(WT)=82, n(2KO)=83; unpaired t-test (\*\*\*\**P* < 0.0001; ns *P* > 0.05). **(c)** Example of primary striatal astrocytes isolated form Pak5/Pak6 dKO mice. Immunocytochemistry was performed with antibodies anti-Arl13b (cilia, green), phalloidin (F-actin, magenta) and with DAPI (blue). Arrows point to primary cilia (scale bar 10 µm). Scale bar of zoomed images is 1 µm. **(d)** Quantification of (c). Violin plots represent the percentage of cells per field with a primary cilium (left) and the cilia length (right). N=3 experiments; n(WT)=64, n(2KO)=52; unpaired t-test (\*\*\*\**P* < 0.0001; ns *P* > 0.05). **(e)** Example of Pak5/Pak6 dKO primary astrocytes transfected with 3xFlag-PAK6 WT or 3xFlag-GUS control. Immunocytochemistry was performed with antibodies anti-Arl13b (cilia, red), Flag (green) and with DAPI (blue). Arrows point to primary cilia (zoomed on the right). Scale bar 10 µm. **(f)** Quantification of (e). Percentage of transfected cells per field with a primary cilium. N=3 experiments; n(GUS)=30, n(PAK6 WT)=28; unpaired t-test (\*\*\**P* < 0.001).

### PAK6 rescues G2019S-but not R1441C-associated ciliogenesis and centrosomal cohesion defects independently from its kinase activity

PAK6 physically and functionally interacts with the PD kinase LRRK2 (33),(34). LRRK2 phosphorylates Rab10 (and other Rabs) at a conserved serine/threonine residue within the switch 2 region and this phosphorylation promotes Rab10 binding to a specific group of interactors, including the ciliary protein RILPL1 (25,42). Mutant LRRK2 hyperphosphorylation of Rab10 at the centrosome/ciliary base results in ciliogenesis and centrosomal cohesion abnormalities (25,27,30,42). Based on (i) our previous observations of a protective action of PAK6 toward mutant LRRK2 (34), (ii) the ability of PAK6 to reduce Rab10 phosphorylation (35) and (iii) the present data supporting PAK6 as a positive regulator of ciliogenesis (**Figures 2-4**), we next tested whether PAK6 can rescue the ciliogenesis and centrosomal cohesion defects associated with mutant LRRK2. To this end, we isolated primary striatal astrocytes from Lrrk2 G2019S knockin (KI) mice and first confirmed previous observations that pharmacological inhibition of LRRK2 kinase activity with MLi-2 is sufficient to rescue G2019S LRRK2-associated ciliogenesis defects (**Figure S1**). Subsequently, G2019S Lrrk2 KI astrocytes were transfected with 3xFlag-PAK6 WT or 3xFlag-PAK6 KM kinase dead along with the 3xFlag-GUS control and the number of ciliated cells that received the plasmid was quantified. Remarkably, both active and inactive PAK6 significantly increased the proportion of ciliated cells (GUS 22%; PAK6 WT 51%; PAK6 KM 44%), indicating that PAK6 restores normal ciliogenesis in G2019S LRRK2 astrocytes and that this ability is independent of its kinase activity (**Figure 5a-b**).

**Figure 5.**
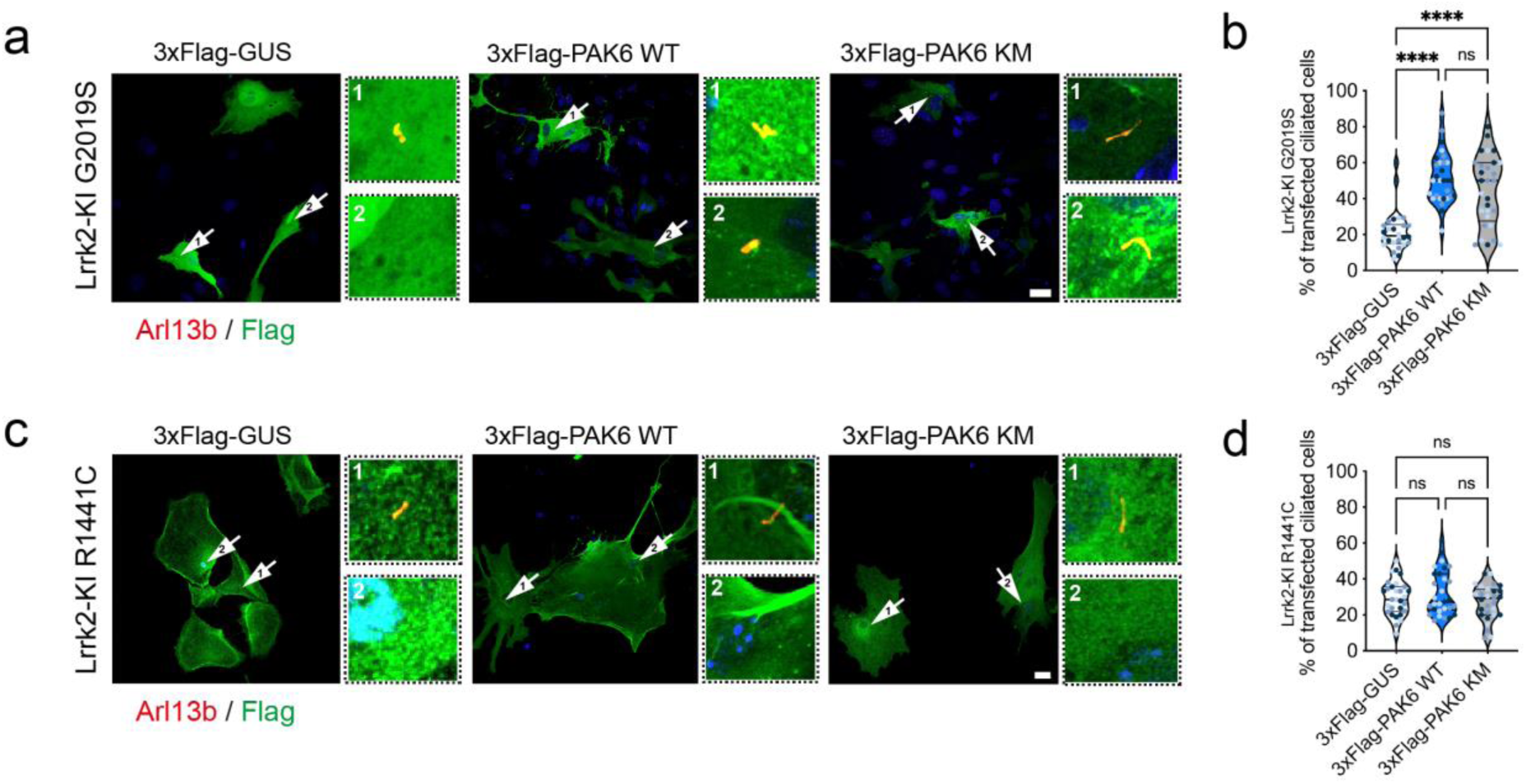
PAK6 rescues G2019S but not R1441C LRRK2-associated ciliogenesis defects independently from its kinase activity. **(a)** Example of primary astrocytes isolated from G2019S LRRK2 KI mice transfected with control 3xFlag-GUS, 3xFlag-PAK6 WT or 3xFlag-PAK6-KM and stained with antibodies against Flag (green), Arl13b (red) and with DAPI (blue). Scale bar 10 µm. Zoomed insets show representative ciliated or non-ciliated cells. **(b)** Quantification of (a). Violin plots represent the percentage of transfected cells per field with a primary cilium. N=3 experiments; n(GUS)=30, n(PAK6 WT)=30, n(PAK6 KM)= 30 fields analyzed; one-way ANOVA with Tukey’s post-test (\*\*\*\**P* < 0.0001). **(c)** Example of primary astrocytes isolated from R1441C KI mice transfected as in (a). Scale bar 10 µm. Zoomed insets show representative ciliated or non-ciliated cells. **(b)** Quantification of (c). N=3 experiments; n(GUS)=29, n(PAK6 WT)=29, n(PAK6 KM)= 28 fields analyzed; one-way ANOVA with Tukey’s post-test (ns *P* > 0.05).

Similar to G2019S LRRK2, the R1441C mutation in the Roc domain of LRRK2 has been linked to defective ciliogenesis (25). To investigate whether PAK6 can also alleviate the ciliogenesis phenotype induced by the R1441C mutant, R1441C Lrrk2 KI primary astrocytes were transfected with 3xFlag-PAK6 WT or 3xFlag-PAK6 KM plasmids and ciliated cells quantified as previously described. Strikingly, neither PAK6 WT nor the kinase dead enzyme were able to modify the proportion of ciliated R1441C LRRK2 astrocytes (**Figure 5c-d**). These results suggest that PAK6 may confer protection only in the presence of the G2019S but not R1441C LRRK2 mutant.

To further test this hypothesis, we used a correlated readout, namely centrosome cohesion deficits that we have previously reported in the presence of mutant LRRK2 (29,30). The proportion of split centrosomes was analyzed in A549 cells transfected with PAK6 WT or PAK6 KM alone or in combination with WT LRRK2, G2019S LRRK2 or R1441C LRRK2. About 20% of A549 cells presented with split centrosomes (**Figure S2a**) and this proportion remained unaltered upon overexpression of PAK6 (WT or KM) alone (**Figure S2a**) or in the presence of WT LRRK2 (**Figure S2b**). When G2019S LRRK2 was overexpressed, the proportion of split centrosomes significantly increased, returning to normal levels upon MLi-2 treatment (**Figure 6a-b**) as previously reported (29,30). The number of split centrosomes also returned to basal levels (control or MLi-2) in cells co-expressing PAK6 WT or KM (**Figure 6a-b**), supporting PAK6 as a modifier of G2019S LRRK2 phenotypes. Conversely, the centrosomal cohesion deficits induced by R1441C LRRK2 were reverted by MLi-2 but not by PAK6 WT or KM expression (**Figure 6c-d**). Thus, PAK6 rescues ciliogenesis and centrosomal cohesion deficits in G2019S but not in R1441C LRRK2-expressing cells through a kinase-independent mechanism.

**Figure 6.**
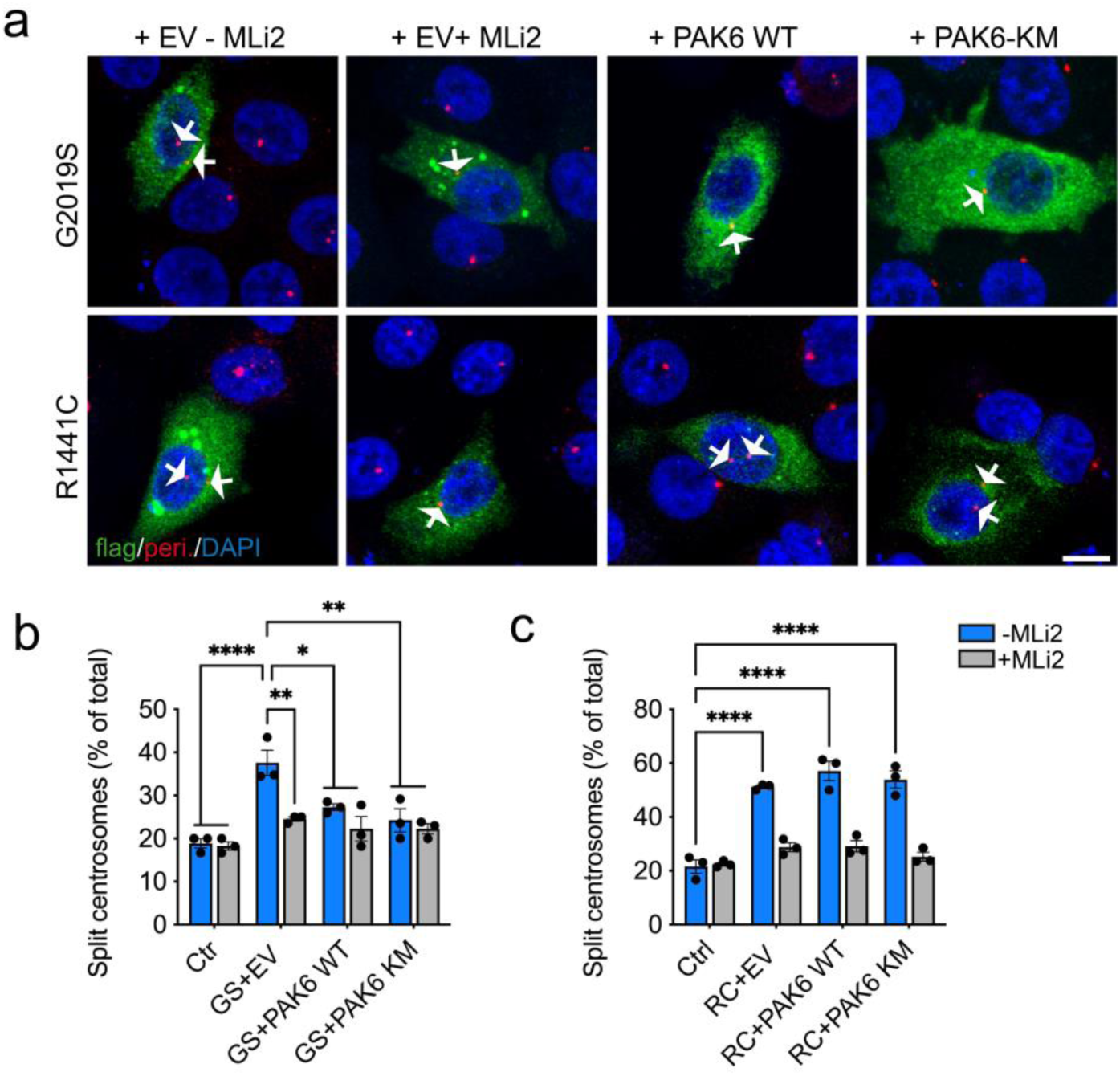
PAK6 rescues G2019S but not R1441C LRRK2-associated centrosomal cohesion defects independently from its kinase activity. **(a)** Example of A549 cells co-transfected with pCMV (EV) and flag-tagged G2019S LRRK2 or R1441C LRRK2 and treated ± MLi-2 (200 nM, 2 h), or co-transfected with flag-tagged LRRK2 and myc-tagged PAK6 WT or PAK6-KM as indicated before immunocytochemistry with antibodies against flag (green), pericentrin (red) and with DAPI (blue). Arrows point to centrosomes in transfected cells. Scale bar, 10 µm. **(b)** Quantification of the percentage of non-transfected cells (ctrl), or cells co-transfected with G2019S LRRK2 and pCMV (EV), PAK6 WT or PAK6-KM and -/+ MLi-2 treatment as indicated where duplicated centrosomes are > 2.5 µm apart (split centrosomes). Bars represent mean ± s.e.m. (n=3 experiments; *p < 0.05; ** p < 0.01; **** p < 0.0001). **(c)** As in (b), but cells co-transfected with R1441C LRRK2. Bars represent mean ± s.e.m. (n=3 experiments; **** p < 0.0001).

### PAK6 affinity to LRRK2 Roc-COR carrying R1441C and Y1699C mutations is dramatically decreased

PAK6 was initially identified to physically bind to the LRRK2 Roc domain through its N-terminal CRIB motif (33). While the G2019S LRRK2 mutation is located in the kinase domain, the R1441C mutation sits in the Roc domain of LRRK2. Thus, we reasoned that one possible mechanism underlying the selective effect of PAK6 toward G2019S but not R1441C LRRK2 could be attributed to a reduced binding with mutant Roc-COR. To test this hypothesis, we evaluated the affinity of recombinant Roc-COR R1441C LRRK2 with recombinant full-length PAK6 using microscale thermophoresis (MST), as we did previously for Roc-COR WT LRRK2 (35). Alongside we also tested the PD mutation Y1699C in the COR domain of LRRK2, which is localized at the Roc-COR interface and nearby the R1441 residue (43). While the affinity of the WT Roc-COR:PAK6 complex is about 10 µM (35), the R1441C mutation decreased the complex K_D_ by 5 times and the Y1699C mutation by 4 times (**Figure 7a**), supporting the notion that Roc-COR mutations weaken PAK6:LRRK2 complex formation. Since PAK6 binds Roc-COR via CRIB, we next used pull-down assays and MST to evaluate the affinity between Roc-COR and CRIB. PAK5 was used as negative control since it does not bind LRRK2 (33) and PAK6 re-expression alone was sufficient to revert the ciliogenesis deficits in dKO Pak5/Pak6 astrocytes (**Figure 4e-f**). PAK5 and PAK6 CRIB domains display a 65% amino acid identity, suggesting some binding specificity toward their PPIs (**Figure 7b**). Pull-down assays confirmed binding of CRIB-PAK6 to Roc-COR in the presence of both GDP and non-hydrolyzable GppNHp, whilst CRIB-PAK5 interaction was barely detectable (**Figure 7c**). These findings were quantitatively corroborated by MST assays: the CRIB-PAK6:Roc-COR complex affinity was around 10 µM, in agreement with the K_D_ of the Roc-COR:PAK6 complex (**Figure 7a**), while the K_D_ of CRIB-PAK5:Roc-COR was 10 times higher, supporting PAK6 as a selective LRRK2 interactor over PAK5 (**Figure 7d**). We complemented these data with AlphaFold2 modelling, which revealed that the PAK6-CRIB domain makes contacts with the Roc-COR interface. Notably, the Arg1441 residue, highlighted in red, is in close proximity to the CRIB binding site, and Tyr1699 interacts with Phe18 and His20 of CRIB (**Figure 7e**). Hence, PAK6 counteracts the ciliogenesis and centrosomal cohesion defects of mutant LRRK2 through an inhibitory binding of CRIB to G2019S LRRK2, but not R1441C (and possibly Y1699C) LRRK2. One consequence could be that this binding antagonizes the access of mutant LRRK2 to its ciliary substrate Rab10. To test this possibility, A549 cells were co-transfected with PAK6 (WT or KM) and G2019S or R1441C LRRK2, and the number of cells with pRab10 and γ-tubulin co-localization was calculated. While MLi-2 treatment completely eliminated phospho-Rab10, PAK6 (WT and KM) expression reduced, but not abolished, the number of LRRK2-expressing cells where pRab10 colocalized with γ-tubulin (**Figure S3a-c**). Conversely, in R1441C LRRK2 expressing cells, the number of cells where pRab10 co-localized with γ-tubulin remained unchanged, in agreement with previous findings (**Figure S3b-d**). We conclude that the protective mechanism of PAK6-CRIB against the G2019S LRRK2-mediated ciliogenesis/centrosomal cohesion deficits depends on CRIB:Roc-COR complex formation and is in part attributable to a negative regulation of localized Rab10 phosphorylation.

**Figure 7.**
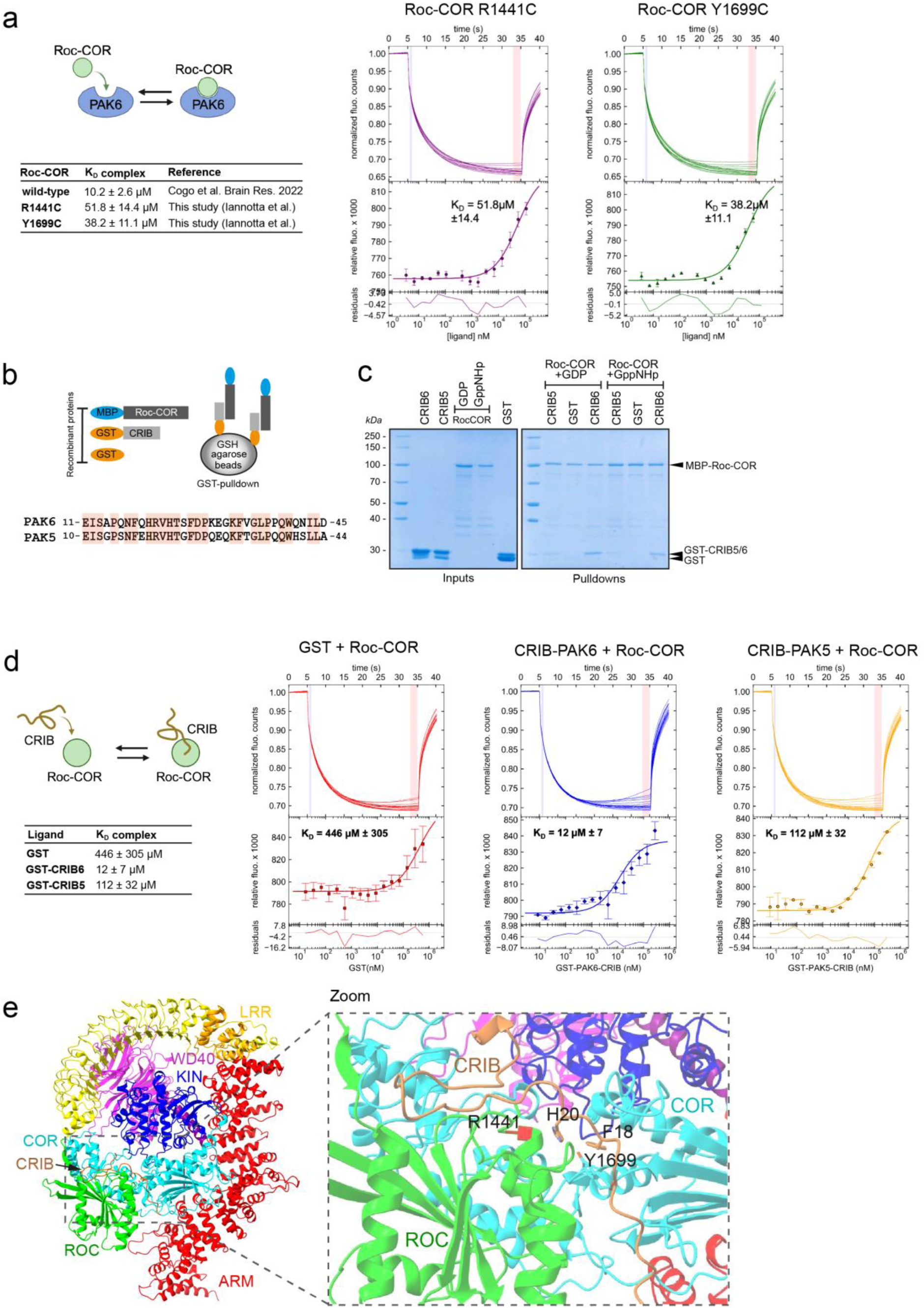
PAK6 affinity to ROC-COR carrying R1441C and Y1699C mutations is dramatically decreased. **(a)** Microscale thermophoresis of fluorescently labelled PAK6 full-length (100 nM) against MBP-fused LRRK2 Roc-COR domain was measured for R1441C (3.4 nM to 110 µM) and Y1699C (3.6 nM to 118 µM) mutations and compared with WT Roc-COR from identical experimental setting published in (35). Upper panel: the normalized fluorescence intensities of all MST traces are plotted against different concentrations of MBP-LRRK2-Roc-COR. Lower panel: The changes of relative fluorescence during thermophoresis. The calculated KD values of PAK6 towards LRRK2-Roc-COR WT, R1441C and Y1699C are 10.2 µM ± 2.6, 51.8 µM ± 14.4 and 38.2 µM ± 11.1 respectively. Error bars show the S.D. of three measurements. The residuals between the data and the fit are shown at the bottom of the graph. **(b)** Schematic of GST-pull down (top) and alignment of PAK5 and PAK6 CRIB domains (bottom), which show 65% of amino acid identity. **(c)** Pull downs of Roc-COR and PAK5/PAK6 CRIB domains. Left panel: Coomassie gel of inputs MBP-Roc-COR purified with Amylose resin and GST, GST-CRIB-PAK5 or GST-CRIB-PAK6 purified with GSH resin. Right panel: pulldowns of GST proteins after incubation with amylose-beads bound Roc-COR in the presence of GDP or non-hydrolysable GppNHp. Roc-COR pulls down GST-CRIB-PAK6, very little GST-CRIB-PAK5 and no GST alone. **(d)** Microscale thermophoresis of fluorescently labelled Roc-COR against GST, GST-CRIB-PAK5 and GST-CRIB-PAK6. Upper panel: the normalized fluorescence intensities of all MST traces are plotted against different concentrations of GST proteins. Lower panel: The changes of relative fluorescence during thermophoresis. The calculated K_D_ values of Roc-COR towards GST, GST-CRIB-PAK6 and GST-CRIB-PAK5 are 446 µM ± 305, 12 µM ± 7 and 112 µM ± 32 respectively. Error bars show the S.D. of three measurements. The residuals between the data and the fit are shown at the bottom of the graph. **(e)** Left. Model obtained out of Alphafold2 modelling for LRRK2 and PAK6, colored as in (43). Only the CRIB domain of PAK6 is shown. Right. Zoomed view on the CRIB domain of PAK6, showing interactions with ROC (green) and COR (cyan) domains of LRRK2. R1441 is highlighted in red, in close proximity with the predicted CRIB binding site. Interaction of F18 and H20 with Y1699 is highlighted.

## Discussion

In this study we showed that the brain enriched kinase PAK6 promotes ciliogenesis by binding a coordinated set of ciliary proteins. PD-linked mutations in the kinase LRRK2 cause neurite shortening (34,44), endo-lysosomal pathway impairment (20,21,45), and ciliogenesis and centrosomal cohesion deficits through hyper-phosphorylation of Rab10 and Rab8a (24,27,29,30). We previously discovered that LRRK2 and PAK6 interact to promote neurite outgrowth (33) through a mechanism involving PAK6-mediated phosphorylation of 14-3-3γ (34). Here we demonstrated in primary neurons and astrocytes as well as in A549 cells that PAK6 expression reverts the ciliogenesis/centrosomal cohesion deficits mediated by G2019S LRRK2 mutation in the kinase domain but not when the GTPase/Roc-COR mutation R1441C was expressed. The mechanism involves a physical binding of the two kinases, which occurs between the N-terminal region of PAK6, termed CRIB (Cdc42 and Rac interactive binding) domain, and the Roc-COR domain of LRRK2 (33). Combining biophysical measurements with recent structural information of full-length LRRK2 (43) coupled to AlphaFold2 modelling (46), we mapped the interaction of CRIB at the Roc-COR interface, providing the framework to interpret the decreased binding between PAK6 and mutant Roc-COR (R1441C and Y1699C). Substitution of tyrosine 1699 with a cysteine is predicted to disturb the interaction with the aromatic ring of P18 and with H20 of CRIB. Similarly, a C1441 mutant should interfere with the correct orientation of H20 side chain, resulting in a weaker binding. In support of the lower affinity of CRIB PAK5 toward Roc-COR, Q19 in CRIB PAK6 is a substituted with a glutamate (E18) in CRIB PAK5 and, according to the model, this substitution is predicted to be unfavorable as it is located in a region of LRRK2 containing two negatively charged aminoacidics, D1799 and E1803, which would cause a repulsion.

CRIB domains are found in different classes of proteins, including p21-activated kinases (PAKs), where they serve as downstream effectors of the small GTPases Cdc42 and Rac1 (1). Group I PAKs (PAK1, PAK2 and PAK3) are activated by Cdc42/Rac1 binding whilst group II PAKs (PAK4, PAK5 and PAK6) are relocalized to signaling sites rather than activated (3). In the context of PAK6 interaction with LRRK2, CRIB enters a pocket formed by the Roc-COR interface, a region that is structurally distinct from that bound by Cdc42 (2ODB PDB structure). Moreover, the N-terminal region of PAK6 following CRIB and proceeding the kinase domain is unstructured in AlphaFold and may require a partner, such as LRRK2, to properly fold. Thus, we can predict a two-step binding involving CRIB entering the Roc-COR pocket and a subsequent stabilization of the complex through PAK6 folding up on other regions of Roc-COR or additional LRRK2 domains.

Future structural predictions coupled with cryo-EM data of full-length proteins will further clarify the mechanism of LRRK2-PAK6 complex formation.

Since LRRK2 PD mutations confer increased kinase activity, ATP-competitive inhibitors are under clinical investigation and hold high therapeutic expectations (47). However, recent structural investigations revealed that type I LRRK2 inhibitors (those that are currently tested in clinical trials) stabilize LRRK2 in a high-affinity oligomeric conformation bound to microtubules (MT) (48,49). Clearly, this property may confer to inhibitor-bound LRRK2 an unwanted gain-of-function, i.e. interference with MT transport, a process particularly important for long range transport of cargos along the axon. Thus, exploring alternative strategies to correct mutant LRRK2 function is highly desirable and our study provides an exciting alternative route for future therapeutic developments. Small molecules stabilizing the LRRK2:PAK6 interaction may be useful in the frame of a personalized medicine approach, namely for patients with mutations outside of the Roc-COR domain. Moreover, the brain-enriched expression of PAK6 may provide an additional advantage for patient stratification with “brain-first” disease (50).

The mechanism underlying centrosomal cohesion and ciliogenesis deficits in mutant LRRK2 depends on a phospho-RAB8a-RAB10/RILPL1 axis (30,51). Our previous study showing that overexpression of active PAK6 decreases phospho-RAB10 levels in WT and G2019S LRRK2 but not R1441G LRRK2 expressing cells (35) suggested to us that the mechanism underlying the mutant-specific rescue of ciliogenesis/centrosomal cohesion defects by PAK6 could be related to the formation of a pool of LRRK2 trapped away from RAB8/RAB10 through PAK6 physical binding. This hypothesis is supported by the reduced number of cells with co-localizing pRAB10/γ-tubulin that we observed when overexpressing PAK6 in G2019S LRRK2 but not in R1441C LRRK2 transfected cells. The incomplete depletion of phospho-RAB10 may be explained by the mechanism relying on stochiometric subtraction of LRRK2 by PAK6 rather than signaling amplification, leading to a reduction of phospho-RAB10 within a safe level (**Figure S3e**). Future experiments should be designed to evaluate whether this hypothesis can be verified, for example by quantitative determination of RAB10 phosphorylation by mass spectrometry (52) in isolated centrosomes (53).

The novel finding of PAK6 as a positive regulator of ciliogenesis reported here may have cellular effects beyond its interaction with LRRK2. PAK6 was previously reported to co-localize with Eg5 in centrosomes in the context of malignant cell transformation (8). Centrosome abnormalities are linked to genomic instability and are considered one possible cause of cancer progression (54). Accordingly, several lines of evidence indicate that PAK6 is overexpressed in advanced cancers including prostate, colon and breast tumors. Thus, too much PAK6 activity may affect cell cycle and promote cell invasion. Based on our data, PAK6 kinase activity is required to promote ciliogenesis whereas the protective effect toward mutant LRRK2 is kinase activity-independent but rather dependent on its CRIB domain. Thus, increasing PAK6-LRRK2 interaction through stabilizing compounds rather than increasing the amount of PAK6 cellular concentration should be considered as a route for future therapeutic developments in stratified LRRK2-PD patients.

## Materials and methods

### HuProt protein microarray and gene ontology enrichment analysis

HuProt^TM^ Human Proteome Microarray v4.0 was purchased from Cambridge Protein Arrays (Babraham Research Campus, Cambridge, UK) and employed to screen PAK6 interactor candidates following manufacturer’s instructions. In detail, protein microarrays were incubated overnight at 4°C with blocking buffer (2% BSA in PBS+0.05% Tween20 (PBST)). Recombinant PAK6 (ThermoFisher, #PR5307A) was diluted in blocking buffer and incubated with the arrays at room temperature (RT) for 2 hours with gentle rocking. After washes with PBST, microarrays where incubated for 2h at RT in gentle rocking with anti-PAK6 1:500 (Abcam, #ab1544752) diluted in blocking buffer, followed by incubation for 2h at RT with fluorophore conjugated antibodies goat anti-Rabbit (AlexaFluor, Invitrogen, #A11036, 1:1000) and anti-GST-Dylight 650 (Columbia Biosciences, #D10-1310, 1:2000). Subsequently, arrays were washed with PBST and 0.1x PBS and dried before being scanned at 532 nm (for detection of sample interactions) and at 633 nm excitation (for detection of GST proteins spotted on the slides) with a resolution of 10 μm.

### Gene ontology (GO) and gene set enrichment analysis (GSEA)

Gene ontology biological process (GO-BP) enrichment analysis was applied to the datasets obtained from microarray analysis via g:Profiler (https://biit.cs.ut.ee/gprofiler/gost) to explore the biological pathways in which candidate interactors are implicated. Query parameters were set as follows: organism: Homo sapiens (Human); Statistical domain scope: Only annotated genes (only genes with at least one annotation); Significance threshold:. g:SCS threshold. Enriched GO terms with the adjusted p < 0.05 were considered statistically significant. Of note, a term size cutoff was set to increase specificity of the enrichment results. Significant GO terms were then grouped based on semantic similarity.

Gene set enrichment analysis (GSEA) was performed to examine the enrichment of cilia-related proteins in the PAK6 array PPI list and the PAK6 PHM PPI list. The pipeline of GSEA was designed as follows: (1) Primary cilium proteome was defined as the gene list annotated to the GO term “cilium” (GO:0005929); (2) The number of overlapping proteins between PAK6 PPI lists and cilium proteome was counted (the “test_intersection”); (3) 10000 randomly sampled gene lists were generated from the MGI gene annotation (N = 24609, all converted to HUGO gene symbols) at same size of the 2 PAK6 PPIs lists, respectively. The overlap sizes between each random gene list and the cilia proteome were counted (the “ref_intersection”); (4) A significant enrichment of cilium-related proteins was defined when the “test_intersection” > 95% of the “ref_intersection” for each PAK6 PPI list.

### Animal models

C57BL/6J Lrrk2 wild-type (WT), Lrrk2 G2019S knock-in (KI), Lrrk2 R1441C KI and Pak5/6 double KO mice were employed. Lrrk2 G2019S KI mice were obtained from Prof. Michele Morari and Novartis Institutes for BioMedical Research, Novartis Pharma AG (Basel, Switzerland) (55). Lrrk2 R1441C KI (56) were obtained from Dr. Huabin Cai (NIH, Bethesda, USA). Pak5/6 knock-out (KO) mice (B6;129-Pak6^tm1Amin^ Pak5^tm1Amin^/J, JAX stock #015825) (5) and WT littermates were obtained from Jackson Laboratory and housed and bred at the University of Padova. PAK6 null mice (B6;129-Pak6^tm1Amin/J^ , JAX Lab) were housed and bred in a climate-controlled vivarium at Florida Atlantic University (FAU). Genotyping was executed using Phire Tissue Direct PCR Master Mix (Thermo Fisher Scientific) and the following primers: Pak5 WT forward, 5’-GCTTCCTCAGATCCATCCAAGGT-3’; Pak5 KO forward, 5’-CTTCCTGACTAGGGGAGGAGT-3’; Pak5 reverse, 5’-AGATGCATTGAGTGCTGGGGAA-3’; Pak6 WT forward, 5’-TCAGTTATCAGCTCCAACACCCTG -3’; Pak6 KO forward, 5’-GCTACCGGTGGATGTGGAATGTGT-3’; Pak6 reverse, 5’-GAGGAAACCCCAGGTCATATACCT-3’. Housing and handling of mice were done in compliance with national guidelines. All animal procedures were approved by the Ethical Committee of the University of Padova and the Italian Ministry of Health (licenses 1041/2016-PR, 105/2019-PR, 200/2019-PR and 690/2020-PR-D2784.N.QHV), by the Institutional Animal Care and Use Committee (IACUC) of FAU and in compliance with the National Institutes of Health Guidelines for the Care and Use of Laboratory Animals and by the NIH guidelines for the Care and Use of Laboratory Animals, Approval number 463-LNG-2021.

### Cell culture and transfection for ciliogenesis analysis

Primary striatal astrocytes were isolated from C57BL/6J Lrrk2 WT, Lrrk2 G2019S KI, Lrrk2 R1441C KI, Pak5/6 KO and relative littermate WT P0-P2 pups as previously described (45,57) and cultured in Basal Medium Eagle (BME, Biowest), supplemented with 10% Fetal Bovine Serum (FBS) (Corning) and 100 U/mL Penicillin and 100 µg/mL Streptomycin (Life Technologies). Astrocytes plated at a density of 2 × 10^5^ cells on 12 mm glass coverslips (VWR) coated with 0.1 mg/mL poly-L-lysine (PLL) were transfected with 1 μg/well of 3xFlag tagged PAK6 (WT or K436M) (33,34). Cells were fixed 36 h after transfection with 4% paraformaldehyde (PFA) in PBS for 20 min at RT.

Primary cortical neurons were obtained as previously described (58) from Pak5/6 KO and relative WT littermate P0 pups. In details, neurons were plated on PLL-coated glass coverslips at a density of 2 × 10^5^ cells/well in Neurobasal A (Life Technologies) supplemented with 2% B27 Supplements (Life Technologies), 0.5 mM L-glutamine (Life Technologies), 100 U/mL penicillin and 100 μg/mL streptomycin. After 10 days in culture, cells were fixed with 4% PFA in PBS for 20 min at RT.

Human neuroblastoma-derived SH-SY5Y cells (naïve), stable lines overexpressing PAK6 (LV-PAK6) or overexpressing PAK6 + PAK6-shRNA (LV-PAK6 + PAK6 shRNA) were cultured in Dulbecco’s Modified Eagle’s Medium (DMEM, ThermoFisher Scientific) and Ham’s F-12 Nutrient Mixture (F12, ThermoFisher Scientific) 1:1, supplemented with 10% FBS and 100 U/mL Penicillin and 100 µg/mL Streptomycin. SH-SY5Y were plated at a density of 0.7 × 10^5^ cells on 12 mm glass coverslips coated with PLL and fixed 24 h later with 4% PFA in PBS for 20 min at RT. SH-SY5Y naïve cells were transfected with 1 μg/well of 2xMyc-PAK6 WT or K436M (previously generated as described in (34) and Lipofectamine 2000 (Thermo Fisher Scientific) and fixed 24 h later with 4% PFA in PBS for 20 min at RT.

HEK293T and mouse embryonic fibroblast (MEFs) WT and Pak6 null cells were maintained in DMEM supplemented with 10% FBS, 1% glutamine and 1% antibiotic-antimycotic solution. To promote cell ciliogenesis, cells were serum-starved overnight (16 h) by lowering the supplemented FBS to 1%. Lentiviral (LV) downregulation was achieved with Dharmacon GIPZ LV shRNA: non-silencing Control (#RHS4346) and PAK6 Specific (#RHS4430-99880759). Cells plated on coverslips were fixed in methanol at −20°C for 10 min. Fixed cells were permeabilized with 0.1% Triton X-100 for 20 min at RT and then incubated with blocking buffer (5% FBS in PBS 1x) for 1 h at RT prior to incubation with antibodies. In details, primary antibodies were diluted in blocking buffer and incubated overnight at 4°C as follows: anti-Arl13b (Proteintech, #17711-1-AP, 1:2000 for astrocytes and cat# 66739-1-Ig for cell lines), anti-γ-tubulin (Proteintech, cat# 66320-1-Ig, 1:200) anti-FLAG® M2 (Sigma, #F1804, 1:400), anti-pericentrin (Abcam, cat#ab28144, 1:200), anti-c-myc (Roche, #11667149001, 1:200), anti-MAP-2 (H-300) (Santa Cruz Biotechnology, #sc20172, 1:200). Anti-PAK6 polyclonal anti-serum was custom generated against glutathione-S-transferase(GST)-PAK6 (aa 292-400) as previously described (11). After 3×5 min washes with PBS, secondary antibodies goat anti-rabbit Alexa Fluor 488 (Invitrogen #A11034), goat anti-rabbit Alexa Fluor 568 (Invitrogen, #A11036), goat anti-mouse Alexa Fluor 568 (Invitrogen, #A11004) and goat anti-mouse Alexa Fluor 488 (Invitrogen, #A11029) were diluted 1:200 in blocking buffer and incubated for 1 h at RT. Subsequently, Hoechst (Invitrogen, 1:10,000) was used as a nuclear counterstain and Phalloidin-647 Reagent (Abcam, #ab176759, 1:100) was used in some experiments to visualize astrocytes. Coverslips were mounted using Mowiol. Fluorescence images were acquired with: Nikon A1R laser scanning confocal microscope, Zeiss LSM700 laser scanning confocal microscope exploiting a 63X oil immersion objective, Leica SP5 confocal microscope using an HC PL FLUOTAR 40x/0.70 oil objective. Around 20 optical sections of selected areas were acquired with a step size of 0.5 µm, and maximum intensity projections of z-stack images were used to manually count the number of ciliated cells and cilia length projections using Fiji-ImageJ software.

### Stable SH-SY5Y PAK6 overexpressing cells

SH-SY5Y cells purchased from ICLC (cat.# HTL95013) were cultured in a 1:1 mixture of Dulbecco’s modified Eagle’s medium (DMEM, Life Technologies) and F12 medium, supplemented with 10% fetal bovine serum (FBS, Life Technologies). Cell lines were maintained at 37°C in a 5% CO_2_ controlled atmosphere. 0.25% trypsin (Life Technologies), supplemented with 0.53 mM EDTA, was employed to generate subcultures. Stable cell lines overexpressing PAK6 wild-type were generated as described in (33). Briefly, the cDNA sequence encoding PAK6 was cloned into the lentiviral plasmid pCHMWS-MCS-ires-hygro. 50 μg/ml hygromycin was utilized for selection. Downregulation of PAK6 expression was performed utilizing an shRNA against human PAK6 (sh-1944, Sigma). Western blot in SH-SY5Y cell extracts was performed with anti-PAK6 (Abcam, #ab1544752) and anti phospho-PAK4/5/6 (Cell Signalling, #3241) antibodies.

### Cell culture and transfections for centrosomal cohesion analysis

A549 cells were cultured in DMEM containing high glucose without glutamine, and supplemented with 10% FBS, 2 mM L-glutamine, 100 U/ml of penicillin and 100 µg/ml of streptomycin as previously described (30).

For centrosomal cohesion determinations, cells were co-transfected with 1 µg of flag-tagged WT-LRRK2, G2019S-LRRK2 or R1441C-LRRK2, and with 100 ng of pCMV or myc-tagged PAK6-WT or K436M. For PAK6-only expression, cells were transfected with 1 µg of pCMV and 100 ng of PAK6 or PAK6-K436M. The following day, cells were treated with DMSO or 200 nM MLi2 for 2 h and then fixed with 4% paraformaldehyde (PFA) in PBS for 15 min at RT, followed by permeabilization with 0.2 % Triton-X100/PBS for 10 min. Coverslips were incubated in blocking solution (0.5 % BSA (w/v) in 0.2 % Triton-X100/PBS) for 1 h at RT, and incubated with primary antibodies in blocking solution overnight at 4 °C. Primary antibodies included rabbit polyclonal anti-pericentrin (Abcam, #ab4448, 1:1000) and mouse monoclonal anti-FLAG® M2 (Sigma, #F1804, 1:500), and rabbit polyclonal anti-myc (Sigma-Aldrich, #C3956, 1:1000). The following day, coverslips were washed 3 x 10 min with 0.2 % Triton-X100/PBS, followed by incubation with secondary antibodies (1:1000) in wash buffer for 1 h at RT. Secondary antibodies included Alexa488-conjugated goat anti-rabbit (ThermoFisher, #A11008) and Alexa594-conjugated goat anti-mouse (ThermoFisher, #A11005). Coverslips were washed three times in wash buffer, rinsed in PBS and mounted in mounting medium with DAPI (Vector Laboratories, H-1200-10).

For determination of pRab10 colocalization with the centrosome, cells were co-transfected with 1 µg of flag-tagged G2019S-LRRK2 or R1441C-LRRK2 and with 100 ng of GFP, or with 1 µg of flag-tagged G2019S-LRRK2 or R1441C-LRRK2 and with 100 ng of mCherry-tagged PAK6 or PAK6-KM as described above. After fixation with 4% PFA, cells were additionally fixed with methanol at -20 °C for 10 min required for γ-tubulin staining. Permeabilization and staining with primary and secondary antibodies was as described above. Cells co-expressing flag-tagged LRRK2 and GFP were co-stained with mouse anti-γ-tubulin (Abcam, #ab11316, 1:1000), and rabbit anti-pRab10 (Abcam, ab241060, 1:1000), followed by co-staining with Alexa405-coupled goat anti-mouse (ThermoFisher, #A31553, 1:1000) and Alexa649-coupled goat anti-rabbit (ThermoFisher, #A21244, 1:1000) secondary antibodies. Cells co-expressing flag-tagged LRRK2 and mCherry-tagged PAK6 were stained sequentially with chicken anti-mCherry (Sigma, #AB356481, 1:1000) followed by Alexa405-coupled goat anti-chicken (Abcam, #ab176575, 1:1000). Coverslips were then co-stained with mouse anti-γ-tubulin and rabbit anti-pRab10, followed by co-staining with Alexa488-coupled goat anti-mouse (ThermoFisher, #A11001, 1:1000) and Alexa647-coupled goat anti-rabbit (ThermoFisher, #A21244, 1:1000).

Images were acquired on an Olympus FV1000 Fluoview confocal microscope using a 60×1.2 NA water objective lens. Images were collected using single excitation for each wavelength separately and dependent on secondary antibodies. Around 10-15 optical sections of selected areas were acquired with a step size of 0.5 µm, and maximum intensity projections of z-stack images analyzed using Fiji software. For each condition and experiment, distances between duplicated centrosomes were quantified from 50-60 transfected cells, with mitotic cells excluded from the analysis. As previously described for A549 cells (30), duplicated centrosomes were scored as split when the distance between their centers was > 2.5 µm. For determination of co-localization of pRab10 with the centrosomal marker γ-tubulin, images were acquired as described above, and co-localization determined from 50-60 transfected cells.

### Sucrose-gradient centrifugation

HEK293 cell extracts were prepared in HEPES buffer (50 mM HEPES, pH 7.4, 150 mM NaCl, 1 mM MgCl_2_, 1 mM EGTA, 0.5% Igepal CA-630, 1 mM dithiothreitol [DTT], and protease inhibitor cocktail) supplemented with 0.1 mM GTP and then clarified by centrifugation (20,000×g for 15 min). The extracts were loaded onto detergent-free 40-60% sucrose gradients and centrifuged at 200,000×g (TLS-55 rotor) for 3 hours. After centrifugation, the gradient fractions were collected and analyzed by western blot with anti-PAK6 and γ-tubulin antibodies.

### Cell culture, transfection and PAK6 purification

pcDNA3 carrying 3xFlag tagged PAK6 was transfected using jetPEI (Polyplus transfection) according to manufacturer’s protocol. Forty-eight hours post-transfection, HEK293 cells were collected, washed once with PBS and lysed in lysis buffer containing 20 mM Tris pH 7.5, 150 mM NaCl, 5 mM MgCl_2_, 1 mM EDTA, 1 mM β-glycerophosphate, 2.5 mM sodium pyrophosphate, 1 mM sodium orthovanadate, 0.5% Tween-20 and 1x Roche complete protease inhibitor cocktail (EDTA free) for 45 min. The lysate was cleared by centrifugation at 20,000 x g for 15 min and the supernatant was incubated with Anti-Flag M2 magnetic beads (Sigma) overnight at 4°C with rotation. Afterwards, the beads were washed 10 times in 5 steps of wash buffers containing 20 mM Tris pH 7.5, 5 mM MgCl_2_, 500 mM NaCl and 0.5% Tween 20 (Buffer A) or 20 mM Tris pH 7.5, 5 mM MgCl_2_, 150 mM NaCl and 0.02% Tween 20 (Buffer B). Purified PAK6 was then eluted with in Buffer B supplemented with 150 µg/ml 3xFlag peptides (Sigma).

### MBP fused LRRK2 Roc-COR and GST-CRIB domain purification

MBP-LRRK2-Roc-COR was expressed in *Escherichia coli* BL21(DE3) cells and purified by Dextrin-sepharose HP using MBP buffer consisted of 20 mM HEPES (pH 8), 200 mM NaCl, 10% Glycerol, 10 mM MgCl_2_, 5 mM 2-mercaptoethanol and 0.5 mM GppNHp. The bound proteins were washed using the same buffer supplemented with extra 10 mM MgCl_2_ and 5 mM ATP before elution with MBP-MST buffer consisted of 50 mM HEPES (pH 8), 800 mM NaCl, 25 mM MgCl_2_, 0.25% Tergitol type NP-40, 10 mM D-maltose and 0.5 mM GppNHp.

GST, GST-CRIB(PAK6) and GST-CRIB(PAK5) were expressed in *Escherichia coli* BL21 (DE3) cells and purified by GSH column using a buffer consisting of 50 mM Tris (pH 7.5), 150 mM NaCl, 5% Glycerol, 5 mM MgCl_2_, 5 mM and 3 mM Dithiothreitol (DTT). The bound proteins were washed and eluted using the same buffer supplemented respectively with 0.5 M NaCl and 10 mM GSH.

### Pulldown assay

The sequence of the CRIB PAK6 and PAK6 domains were obtained by synthesis. Four oligonucleotides (SIGMA-Genosis) complementary 2 by 2 and partially overlapping were designed:

PAK6_GST-CRIB-A-For: AATTCAAATGGAGATCTCAGCGCCACAGAACTTCCAGCACCGTGTCCACACCTCCTTC PAK6_GST-CRIB-A-Rev: GGGTCGAAGGAGGTGTGGACACGGTGCTGGAAGTTCTGTGGCGCTGAGATCTCCATTTG PAK6_GST-CRIB-B-For: GACCCCAAAGAAGGCAAGTTTGTGGGCCTCCCCCCACAATGGCAGAACATCCTGGACTGAC PAK6_GST-CRIB-B-Rev: TCGAGTCAGTCCAGGATGTTCTGCCATTGTGGGGGGAGGCCCACAAACTTGCCTTCTTTG PAK5_GST-CRIB-A_For: AATTCAAATGGAAATATCTGGCCCGTCCAACTTTGAACACAGGGTTCATACTGGGTT PAK5_GST-CRIB-A_Rev: GTGGATCAAACCCAGTATGAACCCTGTGTTCAAAGTTGGACGGGCCAGATATTTCCATTTG PAK5_GST-CRIB-B_For: TGATCCACAAGAGCAGAAGTTTACCGGCCTTCCCCAGCAGTGGCACAGCCTGTTAGCATGAC PAK5_GST-CRIB-B_Rev: TCGAGTCATGCTAACAGGCTGTGCCACTGCTGGGGAAGGCCGGTAAACTTCTGCTCTT

The oligonucleotides were phosphorylated and then annealed to obtain the double strand. Finally, they were ligated with the pGEX-4T-2 plasmid (GE Healthcare Life Sciences) previously digested with the restriction enzymes EcoRI and XhoI. The final product was checked by sequencing.

Twenty-five µg of purified MBP-fused (or MBP alone) proteins were incubated with 50 µl of Amylose resin for 1 h at 4°C in rotation. The resin was then washed 3x with a Washing buffer containing 50 mM Tris (pH 7.5), 150 mM NaCl, 5 mM MgCl_2_, 3 mM DDT and 5% Glycerol. Twenty-five µg of purified GST-fused (or GST alone) proteins were added to the resin and the mix was incubated overnight at 4°C in rotation. The next day the resin was washed 3x with Washing buffer and the proteins were denatured using Laemmli buffer. The samples were boiled for 10 min at 95°C degrees and 15 µl were loaded into a 12% polyacrylamide gel. Blue Coomassie was used for staining.

### MicroScale Thermophoresis (MST)

Human PAK6 or MBP-LRRK2-Roc-COR proteins were purified as previously described and labelled with red fluorescent dye NT-647-NHS in the Monolith NT protein labelling kit according to the manufacturer’s protocol. The unreacted dye was removed from labelled proteins by the gravity flow desalting column provided in the kit with the MST buffer. For labelling MBP-LRRK2-Roc-COR, 5 mM MgCl_2_ and 0.5 mM guanine nucleotide were always supplemented during the labelling process. MST were measured by Monolith NT.115 (NanoTemper). Serial dilution of unlabeled ligand proteins were prepared in MST buffer and mixed with NT-647-NHS labelled proteins at a final concentration of 100 nM, with guanine nucleotides at 0.5 mM. The mixtures were incubated on ice for 30 min, centrifuged at 10,000 x g at R.T. for 1 min and loaded into Monolith premium capillaries (NanoTemper). LED laser power was set to reach around 1200 fluorescence counts for fluorescent detection and IR laser power was set at 80% for MST measurements. Data was analyzed by PALMIST (59) and the graphs were created by GUSSI (60).

### Alpha-fold modelling

DeepMind’s advanced machine learning model, AlphaFold2, was used to predict the structures of complexes between human LRRK2 (Uniprot ref ID Q5S007) and PAK6 (Uniprot ref ID Q9NQU5). The code for AlphaFold2 was downloaded from DeepMind’s official GitHub repository (https://github.com/deepmind/alphafold). The computations were performed on workstation with NVIDIA RTX A5000 (24 Gbytes). Each system was equipped with Linux (Ubuntu 20.04), CUDA11, Python 3.8, and TensorFlow 2.3.1.

Sequence alignments were performed with the standard UniRef90 databases. Calculations with AlphaFold2 were conducted using the recommended configurations provided by DeepMind, with structural templates disabled to obtain de novo models. The 25 complexes predicted were benchmarked with the experimental results.

### Statistical analysis

Statistical analyses were performed with GraphPad Prism 10. Data were analyzed by t-test, one-way or two-way ANOVA test followed by Tukey’s post-hoc test. Significance was set at *P* < 0.05. Significance values are indicated in the figure legends.

## Acknowledgments

This work was supported by the Michael J Fox Foundation for Parkinson Research (EG, AK, J-MT, CM and SH), the University of Padova BIRD funding (EG) and the Busch Biomedical Research Grant (SH). We are very grateful to Dr. Mark R. Cookson and Dr. Alice Kaganovich for sharing primary astrocytes isolated from Lrrk2 R1441C KI mice.

---

**Figure S1.**
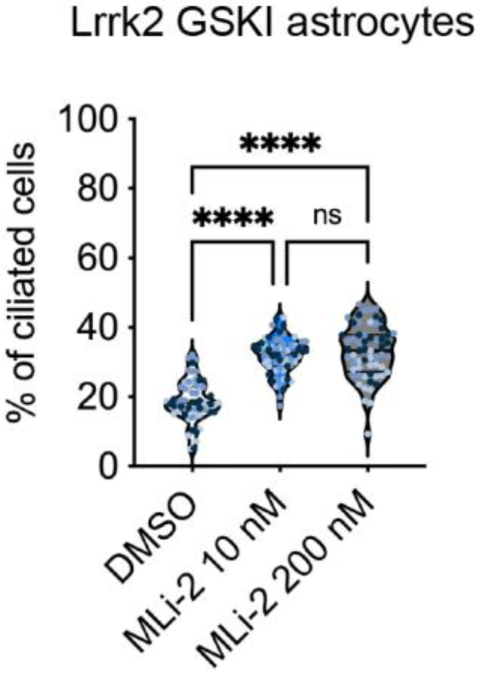
MLi-2 treatment rescues G2019S LRRK2-associated ciliogenesis defects in primary astrocytes. Quantification of the percentage of ciliated G2019S LRRK2 KI primary astrocytes treated with DMSO (n=62), MLi-2 10 nM (n=62) or MLi-2 200 nM (n=62). One-way ANOVA with Tukey’s post-hoc test, \*\*\**P* < 0.001; \*\*\*\**P* < 0.0001.

**Figure S2.**
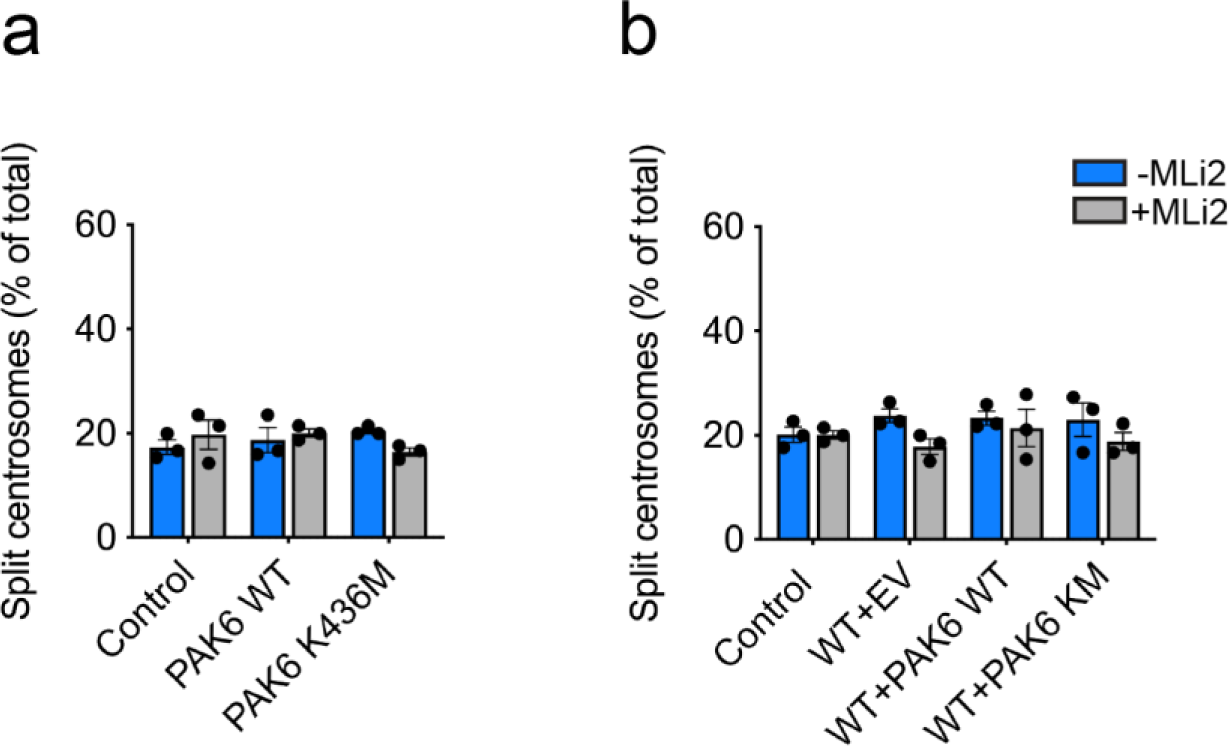
No effect of PAK6 and PAK6-KM on centrosome cohesion in control or wt-LRRK2-expressing cells. **(a)** Quantification of the percentage of non-transfected cells (ctrl), or cells expressing PAK6 or PAK6-KM and -/+ MLi2 treatment as indicated where duplicated centrosomes are > 2.5 µm apart (split centrosomes). Bars represent mean ± s.e.m. (n=3 experiments). **(b)** Quantification of the percentage of non-transfected cells (ctrl), or cells co-transfected with wt-LRRK2 and pCMV (EV), PAK6 or PAK6-KM and -/+ MLi2 treatment as indicated which display duplicated split centrosomes. Bars represent mean ± s.e.m. (n=3 experiments).

**Figure S3.**
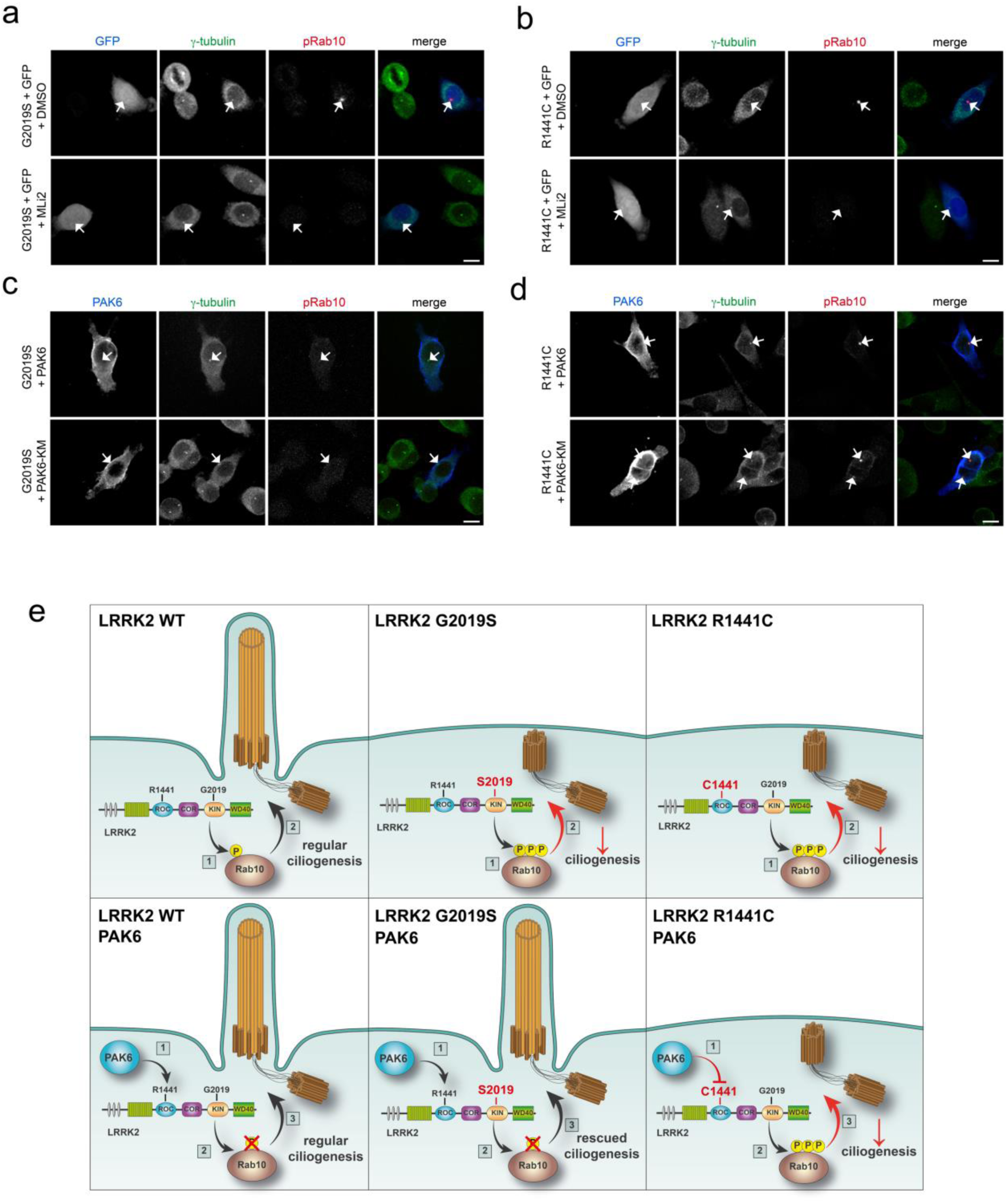
PAK6 partially displaces centrosomal pRab10 in cells expressing G2019S LRRK2 but not R1441C LRRK2. **(a)** Example of A549 cells co-transfected with GFP (pseudo-colored blue) and flag-tagged G2019S LRRK2 and treated with DMSO or MLi-2 (200 nM, 2 h) before staining with γ-tubulin (pseudo-colored green) and pRab10 (far-red). **(b)** Example of A549 cells co-transfected with tagged G2019S LRRK2 and tagged PAK6 or PAK6-KM and stained with antibodies to detect tagged PAK6 (blue), γ-tubulin (green) and pRab10 (far-red). **(c)** Same as (a), but cells co-transfected with GFP and tagged R1441C LRRK2. **(d)** Same as (b), but cells co-transfected with tagged PAK6 or PAK6-KM and tagged R1441C LRRK2. Arrows point to centrosomes in transfected cells. Scale bars, 10 µm. Co-localization of pRab10 and γ-tubulin as quantified from 50-60 transfected cells per condition: G2019S+GFP: 68%, G2019S+GFP+MLi2: 0%; G2019S+PAK6: 55%; G2019S+PAK6-KM: 54%; R1441C+GFP: 78%; R1441C+GFP+MLi2: 0%; R1441C+PAK6: 79%; R1441C+PAK6-KM: 73%. **(e)** Proposed model of PAK6-mediated protection toward LRRK2 G2019S but not R1441C LRRK2 based on this current study and (35).

