## Supplementary figures and images for "PAK6 rescues pathogenic LRRK2-mediated ciliogenesis and centrosomal cohesion defects in a mutation-specific manner"

### Supplementary figure 1

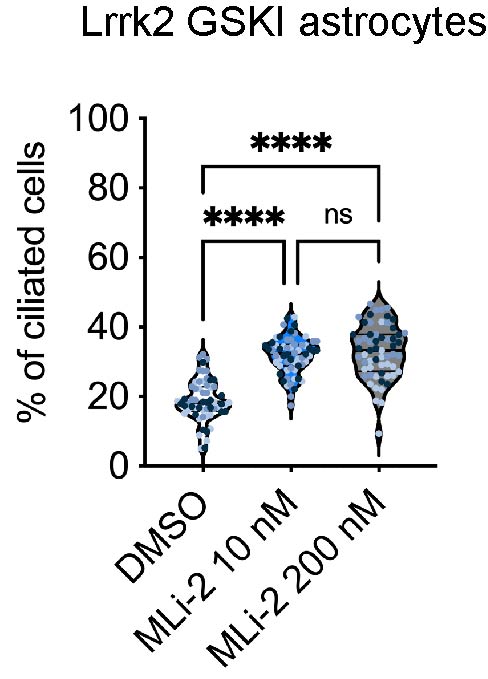

### Supplementary figure 2

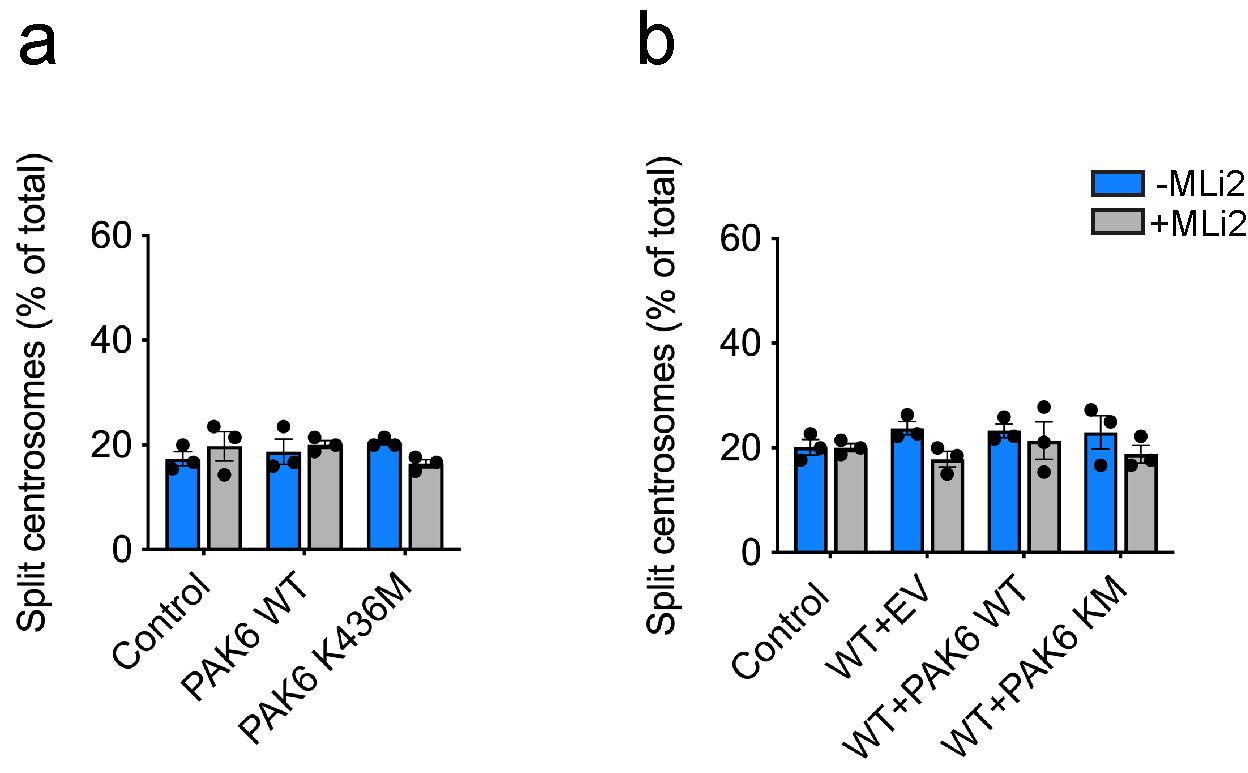

### Supplementary figure 3

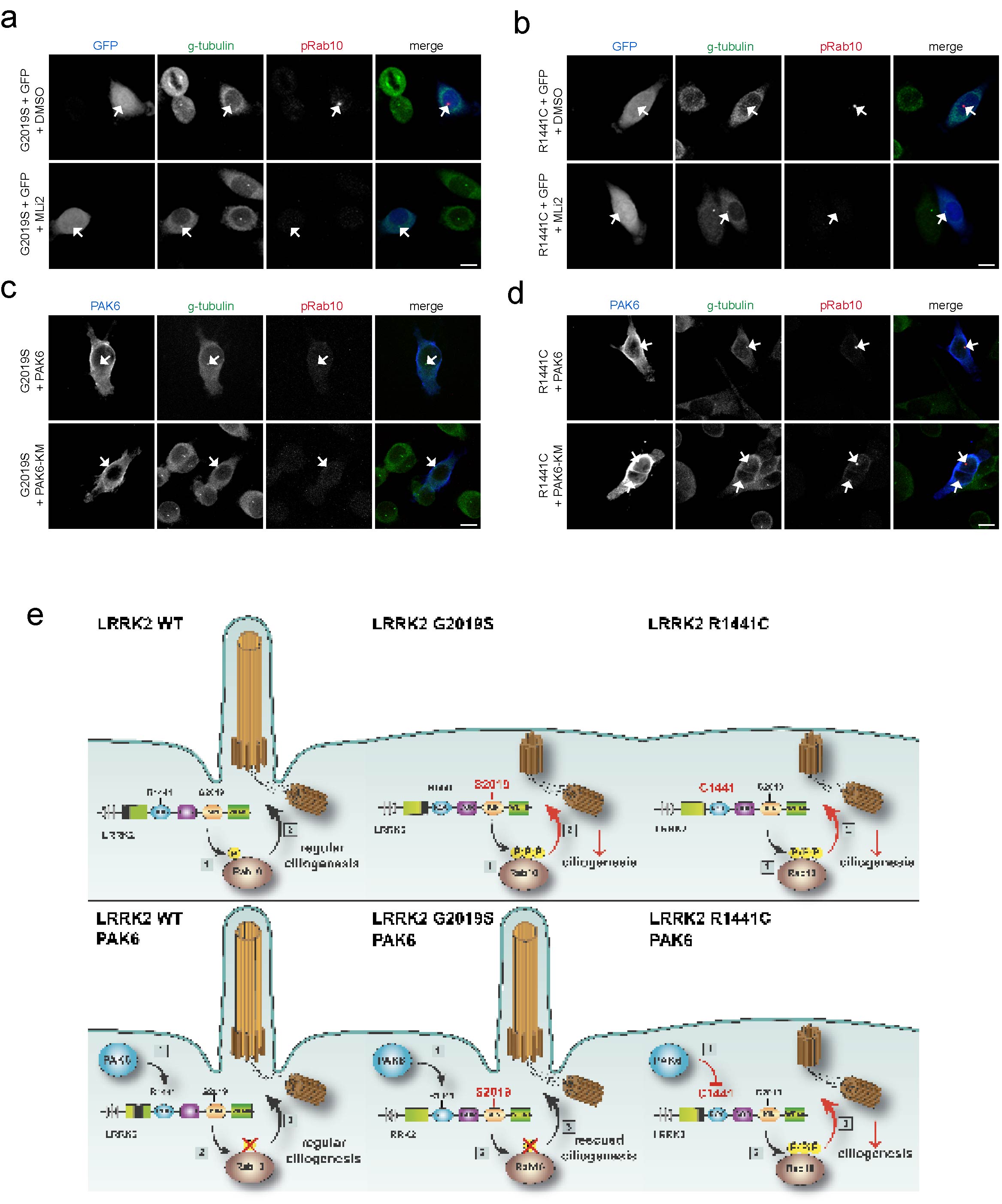
